# The minimal SUF system can substitute for the canonical iron-sulfur cluster biosynthesis systems by using inorganic sulfide as the sulfur source

**DOI:** 10.1101/2024.03.20.586028

**Authors:** Maya Murata, Taichi Murakami, Eiki Yuda, Nanami Mukai, Xintong Zheng, Natsumi Kurachi, Sachiko Mori, Shoko Ogawa, Kouhei Kunichika, Takashi Fujishiro, Kei Wada, Yasuhiro Takahashi

## Abstract

Biosynthesis of iron-sulfur (Fe-S) clusters is indispensable for living cells. Three biosynthesis systems termed NIF, ISC and SUF have been extensively characterized in both bacteria and eukarya. For these L-cysteine is the sulfur source. A bioinformatic survey suggested the presence of a minimal SUF system composed of only two components, SufB* (a putative ancestral form of SufB and SufD) and SufC, in anaerobic archaea and bacteria. Here, we report the successful complementation of an *Escherichia coli* mutant devoid of the usual ISC and SUF systems upon expression of the archaeal *sufB*C* genes. Strikingly, this heterologous complementation occurred under anaerobic conditions only when sulfide was supplemented to the culture media. Mutational analysis and structural predictions suggest that the archaeal SufB*C most likely forms a SufB*_2_C_2_ complex and serves as the scaffold for *de novo* Fe-S cluster assembly using the essential Cys and Glu residues conserved between SufB* and SufB, in conjunction with a His residue shared between SufB* and SufD. We also demonstrate artificial conversion of the SufB*_2_C_2_ structure to the SufBC_2_D type by introducing several mutations to the two copies of *sufB\**. Our study thus elucidates the molecular function of this minimal SUF system and suggests that it is the evolutionary prototype of the canonical SUF system.

## Introduction

Iron-sulfur (Fe-S) clusters are among the most ancient and versatile protein cofactors. They are involved in cellular processes such as respiration, photosynthesis, nitrogen fixation, DNA repair, gene regulation and various other functions throughout all kingdoms of life (Beinert *et al*., 1997; Lill, 2009; Mettert & Kiley, 2015). Although Fe-S clusters are readily inserted into apo-proteins *in vitro* by a simple chemical reaction with iron salts and sulfide, their biosynthesis relies on dedicated systems. Bacteria requires at least one of three distinct biosynthesis systems, NIF, ISC, or SUF. These systems all utilize sulfur mobilized from L-cysteine by the action of cysteine desulfurase (NifS, IscS, or SufS) and assemble a nascent cluster on a so-called scaffold protein (NifU, IscU, or SufBC_2_D) prior to transfer to target apo-proteins (Johnson *et al*., 2005). The NIF system is responsible for the maturation of nitrogenase in diazotrophic bacteria (Dos Santos *et al*., 2007) and is also involved in the maturation of various Fe-S proteins in some non-diazotrophic anaerobes and microaerophiles (e.g.*, Helicobacter pylori*). *Escherichia coli* possesses two biosynthesis pathways: the ISC system in a housekeeping role and the SUF system for backup under stress conditions (Outten *et al*., 2004; Py & Barras, 2010; Takahashi & Tokumoto, 2002; Tokumoto & Takahashi, 2001). These systems are basically redundant in *E. coli* cells except for regulatory mechanisms, and can also be functionally substituted by the foreign NIF system especially under anaerobic conditions (Ali *et al*., 2004; Tokumoto *et al*., 2004).

Despite the difference in the number of the components involved in the NIF and ISC systems, the essential points are shared. NifS and IscS with amino acid identity of more than 30% (Figure S1a) are both classified as Type I cysteine desulfurase (Fujishiro *et al*., 2022; Mihara & Esaki, 2002). The N-terminal domain of NifU is highly homologous to IscU (>30% identity) and functionally essential residues are all conserved (Figure S1b) (Tanaka *et al*., 2019), suggesting that the NIF and ISC systems are mechanistically and evolutionarily related. These characteristics are also seen in the minimal ISC system (also called the MIS system) found in some anaerobes (e.g.*, Clostridium perfringens*) where only two components homologous to IscS and IscU (the second of which we call IscU* for clarity here) are encoded in their genomes (Figure 1) (Garcia *et al*., 2022; Sato *et al*., 2021). Hence, it is likely that auxiliary components of the canonical ISC machinery (Fdx, HscA, HscB and IscA) were recruited to the minimal ISC system during evolution in a process of adaptation to oxygenic environments (Figure 2a). In support of this view, we note that Fdx and IscA in *E. coli* ISC system have been demonstrated to be essential for aerobiosis but not for anaerobiosis (Tanaka *et al*., 2016). Likewise, the function of HscA and HscB could be bypassed well in anaerobiosis by several point mutations in *E. coli* IscU (Sato *et al*., 2021).

**Figure 1.**
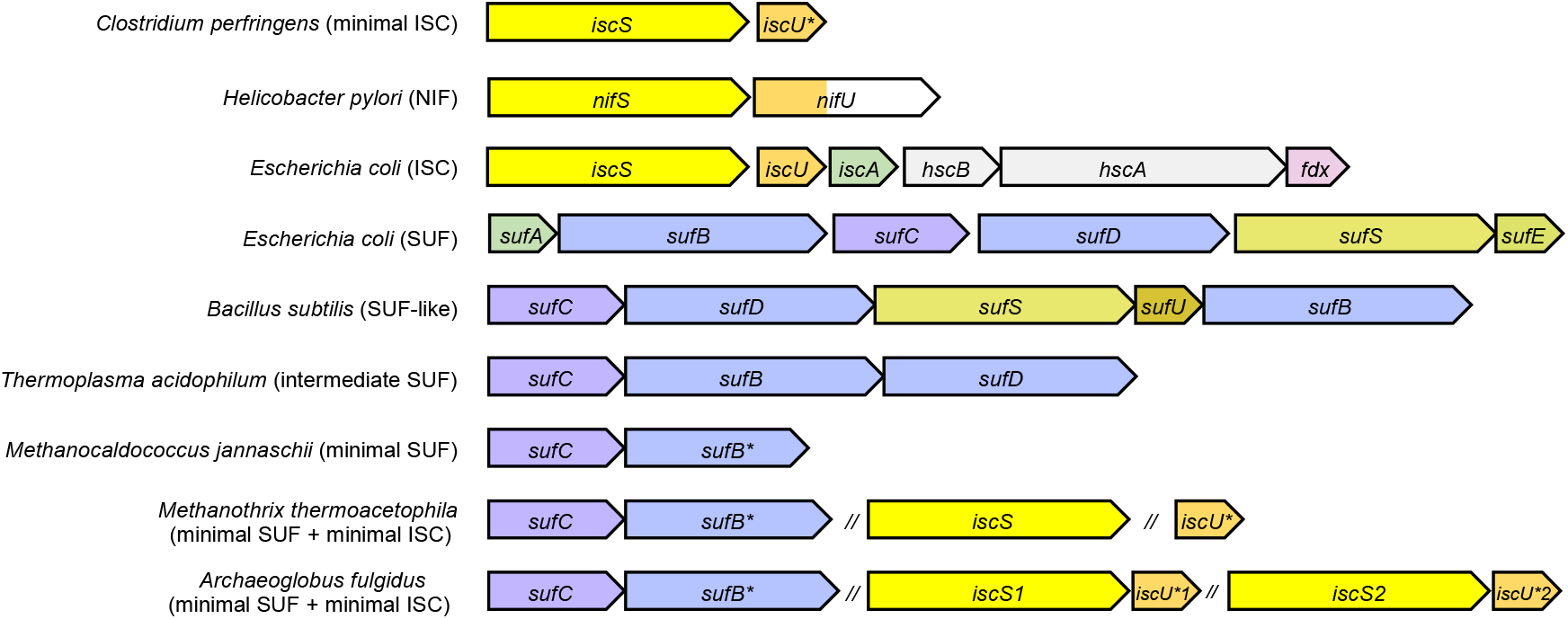
Comparison of gene regions of typical biosynthetic systems for Fe-S clusters in microorganisms. For simplicity, genes for transcription regulators (e.g., *iscR*) are omitted from the figure.

**Figure 2.**
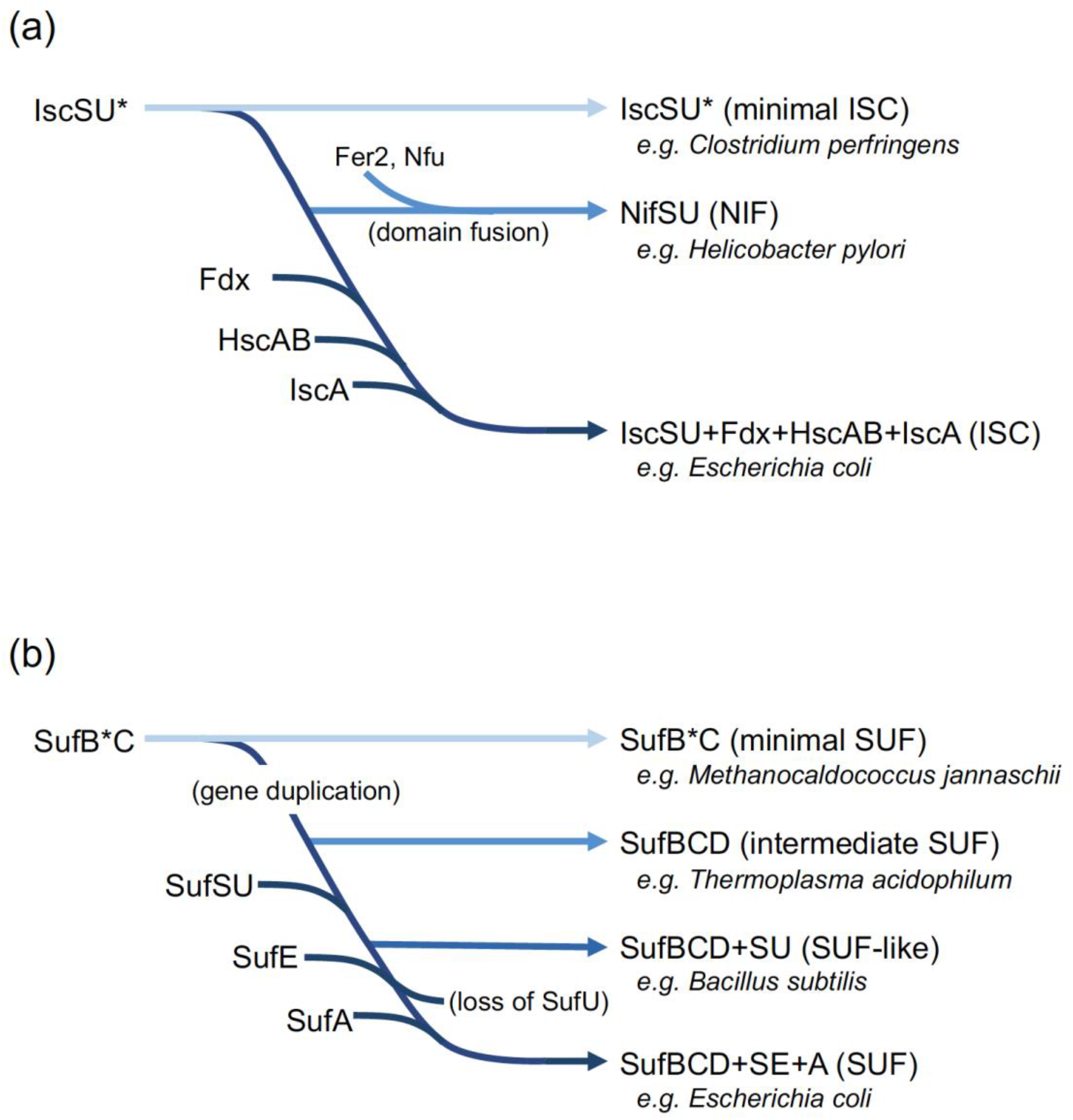
Evolutionary trajectory of the Fe-S cluster biosynthesis systems. (a) The ISC and related systems. (b) SUF and related systems.

In the canonical SUF system in *E. coli*, the sulfur atom is transferred from SufS (Type II cysteine desulfurase) to SufE (sulfur-transfer protein) and then to the SufBCD complex that serves as a scaffold for *de novo* Fe-S cluster assembly (Layer *et al*., 2007; Loiseau *et al*., 2003; Outten *et al*., 2003; Wollers *et al*., 2010). SufA is a so-called A-type carrier protein of the Fe-S cluster (Chahal *et al*., 2009; Gupta *et al*., 2009). As a variation of the SUF system, a number of other bacteria (e.g., Bacilli, Actinobacteria, Thermotogae, and some Spirochaetes) utilize SufU as a sulfur-transfer protein in place of SufE (Fujishiro *et al*., 2017; Selbach *et al*., 2014). Interestingly, SufU exhibits only limited similarity (<30% identity) with IscU of the ISC system (Figure S1b), and evolutionary trajectories of these U-type proteins (IscU*, NifU, IscU and SufU) are inferred to lie elsewhere (Yokoyama *et al*., 2018). As for another variation of the SUF system, its minimal system (also called SMS to indicate that it is the SUF-like minimal system) has been suggested to function in some anaerobic bacteria and archaea (e.g.*, Methanocaldococcus jannaschii*). In this system only two components homologous to SufC and SufB (the second referred to as SufB* here) are encoded in their genomes (Boyd *et al*., 2014; Garcia *et al*., 2022; Tokumoto *et al*., 2004). The SufC sequences are well conserved (>35% identity) between the minimal and canonical SUF systems (Figure S1c), but the SufB* sequences are divergent, with only limited similarity (around 20% identity) with either SufB or SufD. However, the amino acids essential for the function of *E. coli* SufB (Cys254, Cys405, and Glu434) and SufD (His360) (Yuda *et al*., 2017) are strictly conserved in almost all SufB* sequences (Figure S1d). A phylogenetic study indicated that the related sequences can be classified into three subfamilies, SufB*, SufB and SufD, which diverged early during the evolution of life (Figure 3). It has been repeatedly suggested that gene duplication of SufB* and numerous subsequent mutations resulted in SufB and SufD (Boyd *et al*., 2014; Tokumoto *et al*., 2004).

**Figure 3.**
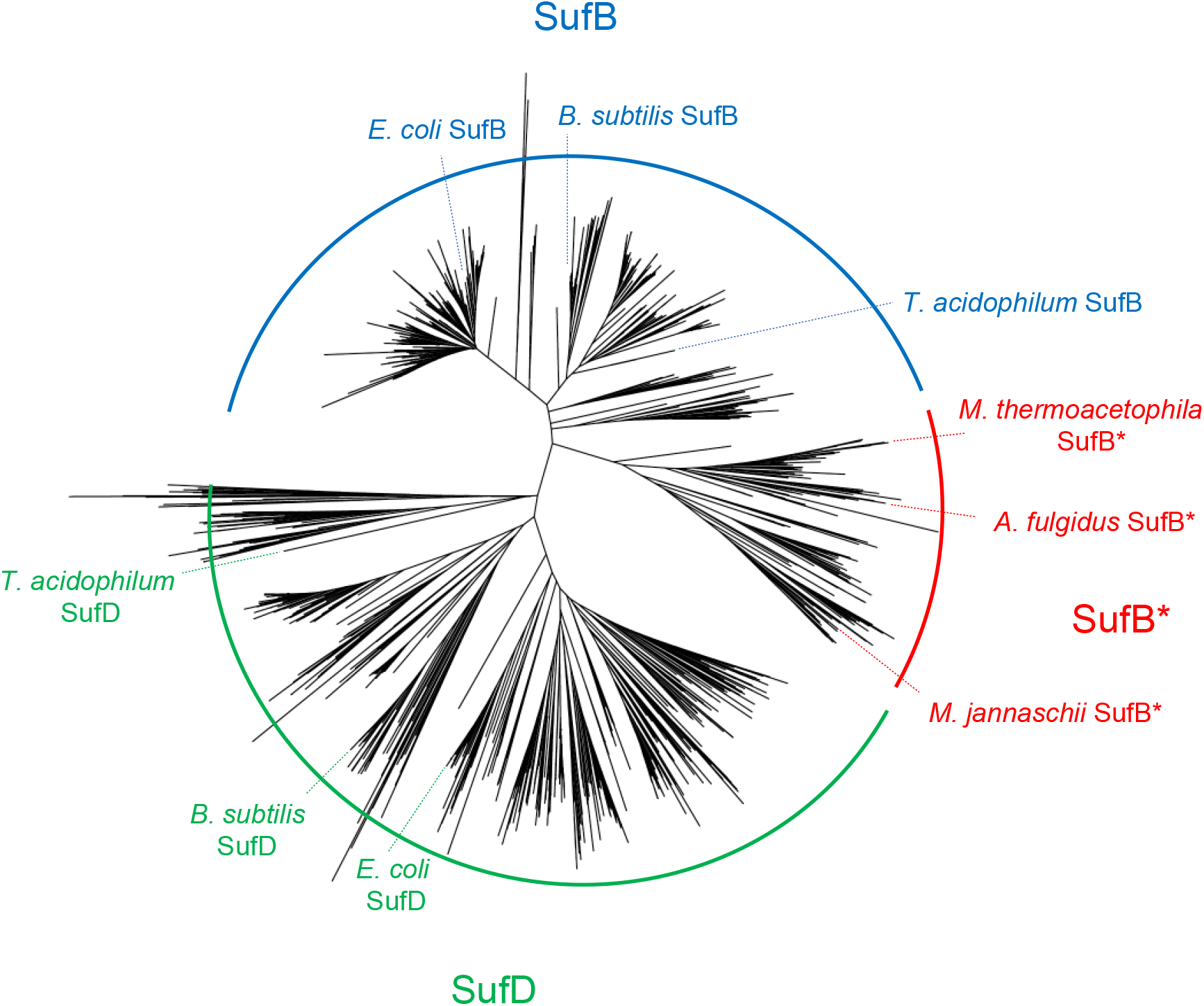
Phylogenetic tree of SufB, SufD, and SufB*. The 1473 amino acid sequences homologous to SufB and SufD were retrieved from a selected subset of the MGBD microbial genome database (http://mbgd.genome.ad.jp) that contained 866 bacterial, 96 archaeal and 48 eukaryotic complete genome sequences. The evolutionary history was inferred using the maximum-likelihood method in the MEGA X software (Kumar *et al*., 2018). The SufB* subfamily is characterized by the absence of the SufD sequence in the respective genomes.

So far, the characterization of the minimal SUF system has been limited. Several lines of genetic evidence indicated that *sufCB\** is essential for viability of the methanogenic archaeon *Methanococcus maripaludis* (Liu *et al*., 2010; Sarmiento *et al*., 2013). By contrast, two copies of *sufCB\** in *Methanosarcina acetivorans* could be both deleted without noticeable defects, possibly because this methanogen contains three additional copies of *iscSU\** (Deere *et al*., 2020; Saini *et al*., 2023). Recently, recombinant SufB*C complexes have been anaerobically purified with Fe-S clusters that could be readily transferred to apo-aconitase *in vitro* (Garcia *et al*., 2022). However, these observations are not sufficient to prove that SufB*C serves as a scaffold for *do novo* assembly of Fe-S clusters. For this study, we examined whether the minimal SUF system can substitute for the role of canonical SUF and ISC systems by taking advantage of an *E. coli* mutant that can conditionally survive without Fe-S clusters, and found that the heterologous complementation occurred under anaerobic conditions when Na_2_S was supplemented to the culture medium but not without supplementation. We also demonstrate that the SufB*C can be converted to the SufBCD type by introduction of several mutations to the two copies of *sufB\**. The evolutionary trajectory of the SUF system will be discussed.

## Results

### *Methanocaldococcus jannaschii* SufB*C can serve as an Fe-S cluster assembly machinery in *E. coli* cells

The coding regions for *sufB\** and *sufC* from the methanogenic archaea *Methanocaldococcus jannaschii* (*Mj*) were synthesized so as to optimize the codons for expression in *E. coli*. These genes were cloned in tandem into the plasmid pBBR-*Mj sufB*C* with artificial ribosome binding sequences, and expressed in *E. coli* UT109 cells, which contains deletions of the chromosomal *isc* (Δ*iscUA-hscBA*) and the *suf* (Δ*sufABCDSE*) operons. Normally, deletion of both pathways in *E. coli* is lethal (Outten *et al*., 2004; Takahashi & Tokumoto, 2002), but the plasmid pUMV22 Sp^r^ carrying genes for mevalonate (MVA) kinase, phosphomevalonate kinase, and diphosphomevalonate decarboxylase for an alternative isoprenoid biosynthesis MVA pathway allows UT109 to grow on rich media with a strict dependency on supplemented MVA (Tanaka *et al*., 2016). Upon introduction of a functional *sufABCDSE* or *iscUA-hscBA*, the cells become able to grow normally even in the absence of MVA. Upon introduction of the plasmid pBBR-*Mj sufB*C*, no growth was observed for the cells under the normal growth conditions lacking MVA. To our surprise, however, heterologous complementation occurred under anaerobic and Na_2_S-supplemented conditions for which both *Mj sufB\** and *Mj sufC* genes were required (Figure 4a). The growth on MVA-free LB glucose media was dependent on the concentration of Na_2_S, which could be as low as 1 mM, whereas the addition of L-cysteine (1 mM) was ineffective (data not shown). Like many other Archaea, *M. jannaschii* possesses the gene for the ApbC/Nbp35 homolog which is presumed to have some kind of (as yet unspecified) role in Fe-S cluster biosynthesis (Boyd *et al*., 2009). However, *Mj apbC* alone was unable to complement the MVA-dependent growth phenotype of UT109 even under anaerobic and Na_2_S-supplemented conditions (Figure S2). When coexpressed with *Mj sufB*C* from compatible plasmids, no stimulatory or inhibitory effects of *Mj apbC* were observed in the heterologous complementation. In another methanogenic archaeon, *M. maripaludis*, the gene for Nbp35/ApbC homolog was disrupted with negligible alteration of the phenotype whereas the *sufB\** and *sufC* were essential for viability (Liu *et al*., 2010; Zhao *et al*., 2020). No further characterization for *Mj* ApbC was undertaken for this study.

**Figure 4.**
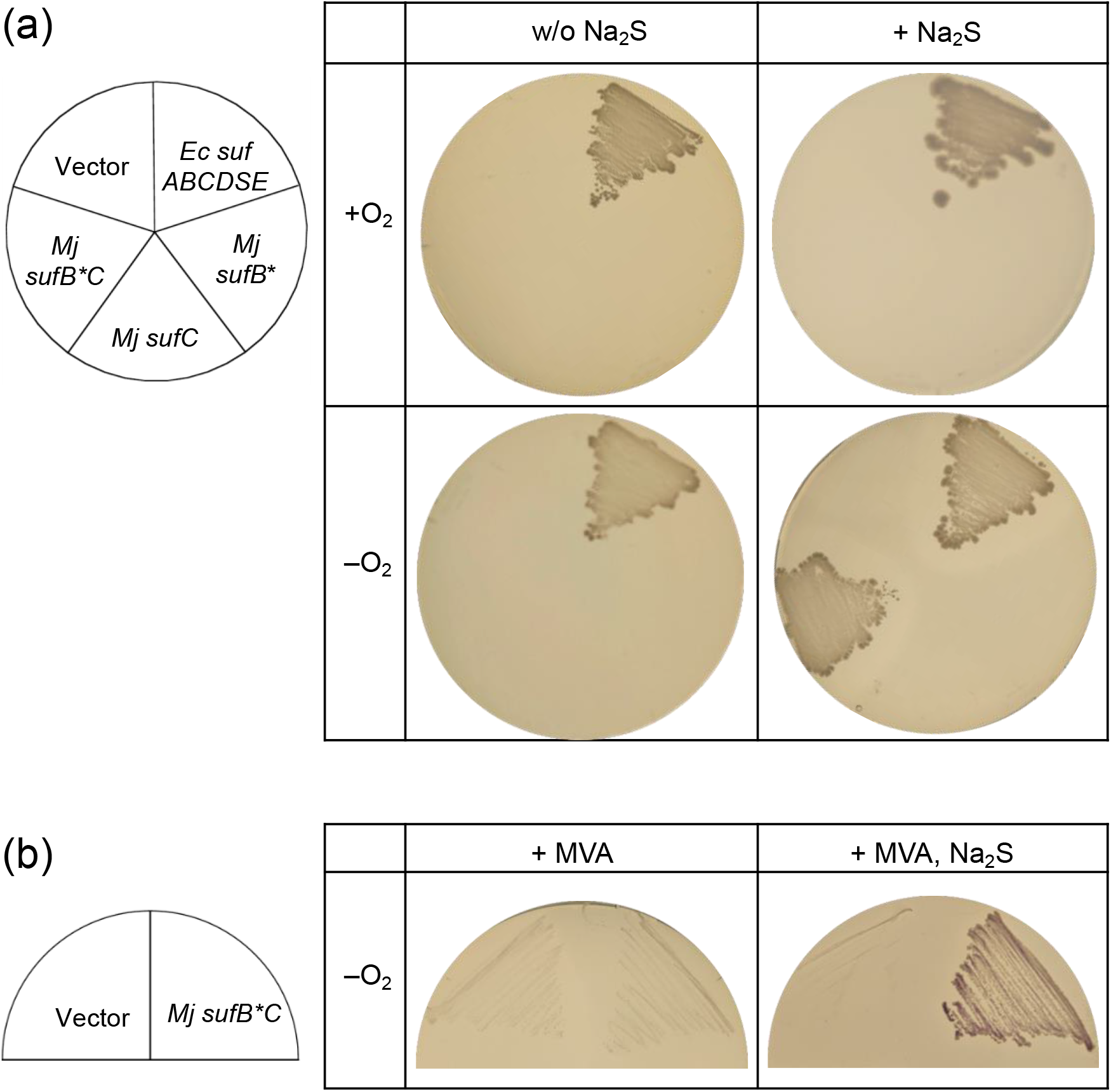
Heterologous complementation of the *E. coli mutant* UT109 by *M. jannaschii sufB*C*. (a) Growth phenotype of the UT109 (Δ*iscUA-hscBA* Δ*sufABCDSE*) cells harboring the plasmid expressing *Mj sufB\** and/or *sufC.* The UT109 cells harboring pUMV22 Sp^r^ were transformed with the pBBR1MCS4-derived plasmids in which *Mj sufB\** and/or *sufC* were cloned. The transformants were once selected on agar plates containing antibiotics and MVA, and then grown on LB-glucose plates (without MVA) at 37°C for 72 hr under aerobic (+O_2_) or anaerobic (–O_2_) conditions. Where indicated, Na_2_S was supplemented at a concentration of 5 mM. For control experiments, the *E. coli suf* operon was expressed from the plasmid pBBR-*sufABCDSE.* (b) Activity staining of the cells for the Fe-S enzyme FDH_H_. Cells were grown anaerobically on LB-glucose plates in the presence of 300 µM MVA with or without 5 mM Na_2_S at 37°C for 24 hr. Each plate was overlaid with a dye solution containing 0.75% agar, 1 mg/ml of benzyl viologen, 0.25 µM sodium formate, and 25 mM KH_2_PO_4_. The color of the cells gradually turned reddish purple with the formate-dependent reduction of benzyl viologen.

The restoration of an MVA-independent growth of UT109 is a clear indication that [4Fe-4S] cluster-containing enzymes IspG and IspH resumed their roles in the isoprenoid biosynthesis MEP pathway (Tanaka *et al*., 2016). To examine Fe-S cluster integration in other Fe-S proteins, the activity of endogenous formate dehydrogenase H (FDH_H_), an anaerobic enzyme containing an oxygen-labile [4Fe-4S] cluster, selenocysteine and molybdopterin guanine dinucleotide was assessed by agar overlay assay. In this experiment, the cells were grown anaerobically on LB agar plates containing MVA and glucose. Upon agar overlay, the cells harboring pBBR-*Mj sufB*C* changed their color to purple due to formate-dependent reduction of benzyl viologen, which was observed only for cells grown in the presence of supplemented Na_2_S (Figure 4b). By contrast, FDH_H_ activity was negligible in the cells grown in the absence of Na_2_S or in the cells harboring the empty vector. These results suggest that when Na_2_S is supplemented to the media, the sulfur moiety permeates through the lipid bilayer probably in the form of H_2_S and is utilized as a sulfur source for the *de novo* assembly of Fe-S clusters that is executed by *Mj* SufB* and SufC.

### Distinct properties of *Mj* SufB*C and *Ec* SufBCD

In the Fe-S cluster biosynthesis systems characterized in research up to now, L-cysteine serves as the universal sulfur source of the cluster. In the *E. coli* SUF machinery, the sulfur is abstracted from L-cysteine by cysteine desulfurase SufS and then delivered to the SufBCD complex in the form of persulfide (-SSH) via the sulfur-transfer protein SufE (Layer *et al*., 2007; Outten *et al*., 2003). In order to examine whether the exogenously added sulfide can also serve as a sulfur source for the *E. coli* SUF machinery, we expressed the *suf* operon lacking the genes for the sulfur supplying system in UT109. When the system was devoid of SufS, SufE, or both, no complementation was observed even under anaerobic and Na_2_S-supplemented conditions (Figure S3) suggesting that, unlike *Mj* SufB*C, the *Ec* SufBC_2_D complex is unable to utilize the sulfur derived from exogenous Na_2_S. We also examined whether the *Ec* SufBC (lacking SufD), *Ec* SufBD (lacking SufC) or *Ec* SufCD (lacking SufB) could complement the phenotype of UT109 under anaerobic and Na_2_S-supplemented conditions, but none of them worked. Taking these results together, we observe that *Mj* SufB* and SufC are unique in that they work as a cluster biosynthesis system with only two components, and that the system uses sulfide instead of L-cysteine. We also examined the combination of *Mj* SufB*C and *Ec* SufASE (lacking SufBCD), but there was no indication that *Mj* SufB*C would receive L-cysteine-derived sulfur from *Ec* SufSE (data not shown).

### Probable architecture of the SufB*C complex

When *Mj sufB*C* was coexpressed in *E. coli*, the harvested cells exhibited a brownish color, but the color was lost during purification under aerobic conditions. Using Ni-NTA chromatography, N-terminal His_6_ tagged *Mj* SufB* was copurified with untagged *Mj* SufC with an approximate 1:1 stoichiometry (Figure S4), suggesting the presence of a SufB*C complex as observed in previous studies (Garcia *et al*., 2022). AlphaFold2 predicted the SufB*_2_C_2_ complex not only for *Mj* SufB*C but also for phylogenetically distant homologs as described below, in which the homodimer interface of SufB*-SufB* consists of two anti-parallel β-sheets (Figure 5). The overall architecture is similar to that of the *Ec* SufBC_2_D complex (Hirabayashi *et al*., 2015) and also consistent with the previous structural data for the *Methanosarcina mazei* Go1 SufB* homodimer (PDB entry 4DN7). Although the two molecules of SufC in the SufB*_2_C_2_ complexes appear to associate with each other in the AlphaFold2 prediction, it should be noted that in the predicted structure of the *Ec* SufBC_2_D complex, the two molecules of SufC are also in close vicinity with each other and in fact closer than in the crystal structure (Figure 5). The exact spatial arrangement of the two molecules of SufC in the actual complex thus needs to be clarified experimentally.

**Figure 5.**
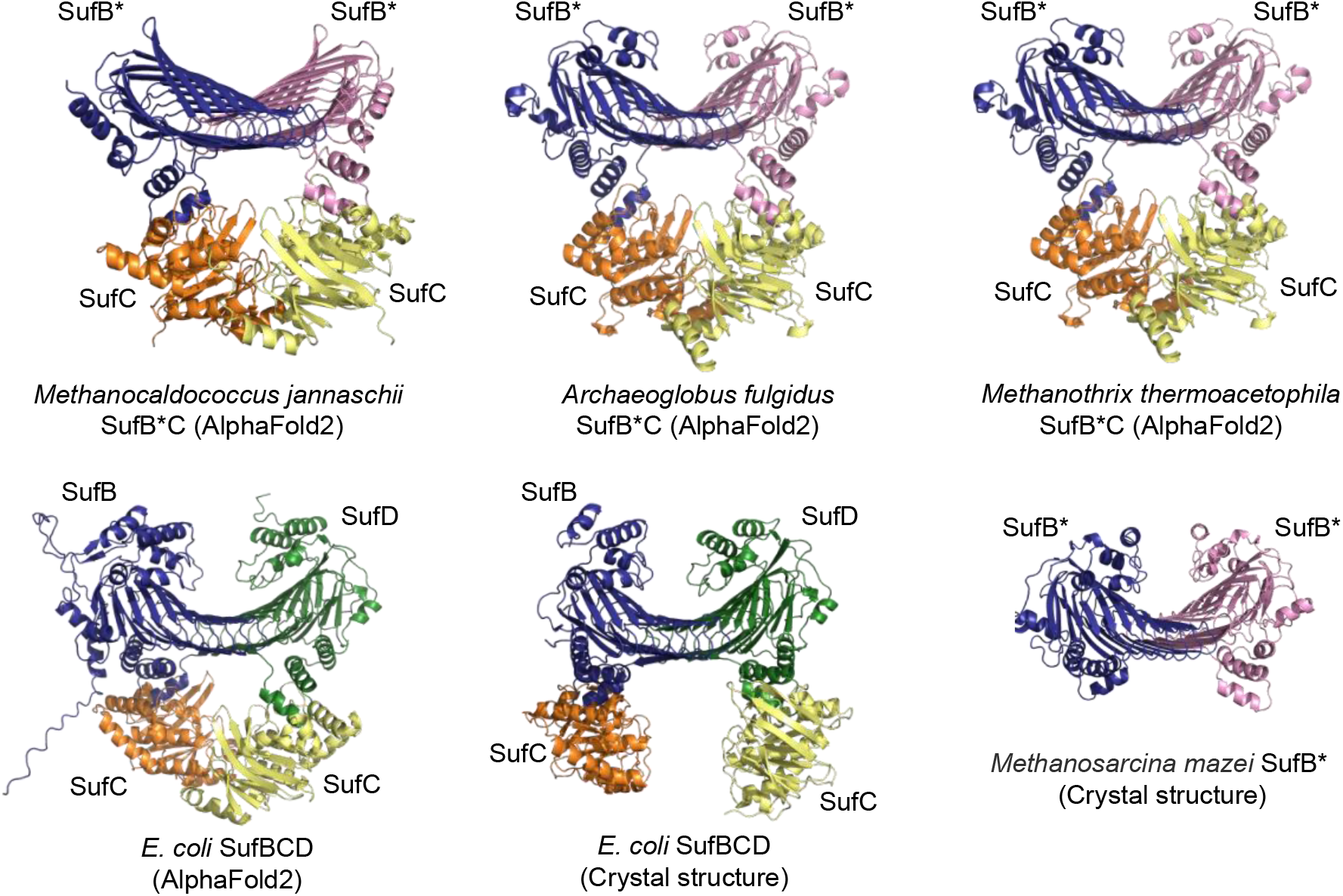
AlphaFold2-predicted structures of the SufB*_2_C_2_ complexes and the SufBC_2_D complex. Structure predictions were performed by using ColabFold v1.5.2-patch: AlphaFold2 using MMseqs2 (Mirdita *et al*., 2022). For comparison, crystal structures of the *E. coli* SufBC_2_D complex (PDB code 5AWF) and *Methanosarcina mazei* SufB*_2_ homodimer (PDB code 4DN7) are included.

### Conservation of functional amino acids between *Mj* SufB*_2_C_2_ and *Ec* SufBC_2_D

We have previously identified amino acids essential for the function of the *Ec* SufBC_2_D complex (Hirabayashi *et al*., 2015; Yuda *et al*., 2017). These residues are conserved in almost all homologous sequences including *Mj* SufB* and SufC. Amino acid substitution of *Mj* SufC K45R (Walker A motif), E171Q (Walker B motif), or H203A (H-motif) abolished heterologous complementation under anaerobic and Na_2_S-supplemented conditions (Figure 6). In *Mj* SufB*, alanine substitution of Cys252, His279, or Glu280 also abolished the complementation of UT109. The corresponding residues are *Ec* SufB Cys405, *Ec* SufD His360, and *Ec* SufB Glu434, respectively, which reside in close vicinity at the interface between *Ec* SufB and SufD and appear to serve as the assembly site for the nascent Fe-S cluster (Hirabayashi *et al*., 2015; Yuda *et al*., 2017). The loss-of-function variants of the *Mj* SufB*C complex were purified by Ni-NTA chromatography as observed for the wild-type complex (Figure S4), confirming that the substitutions in *Mj* SufB* or SufC did not impair the assembly of the *Mj* SufB*_2_C_2_ complex. These results suggest mechanistic similarities between *Mj* SufB_2_*C_2_ and *Ec* SufBC_2_D, in which the interface of the two β-helix core domains (between *Mj* SufB*and SufB* or between *Ec* SufB and SufD) may serve as the assembly site for the nascent Fe-S cluster and dimerization of two SufC molecules coupled with the ATPase activity is required for cluster formation.

**Figure 6.**
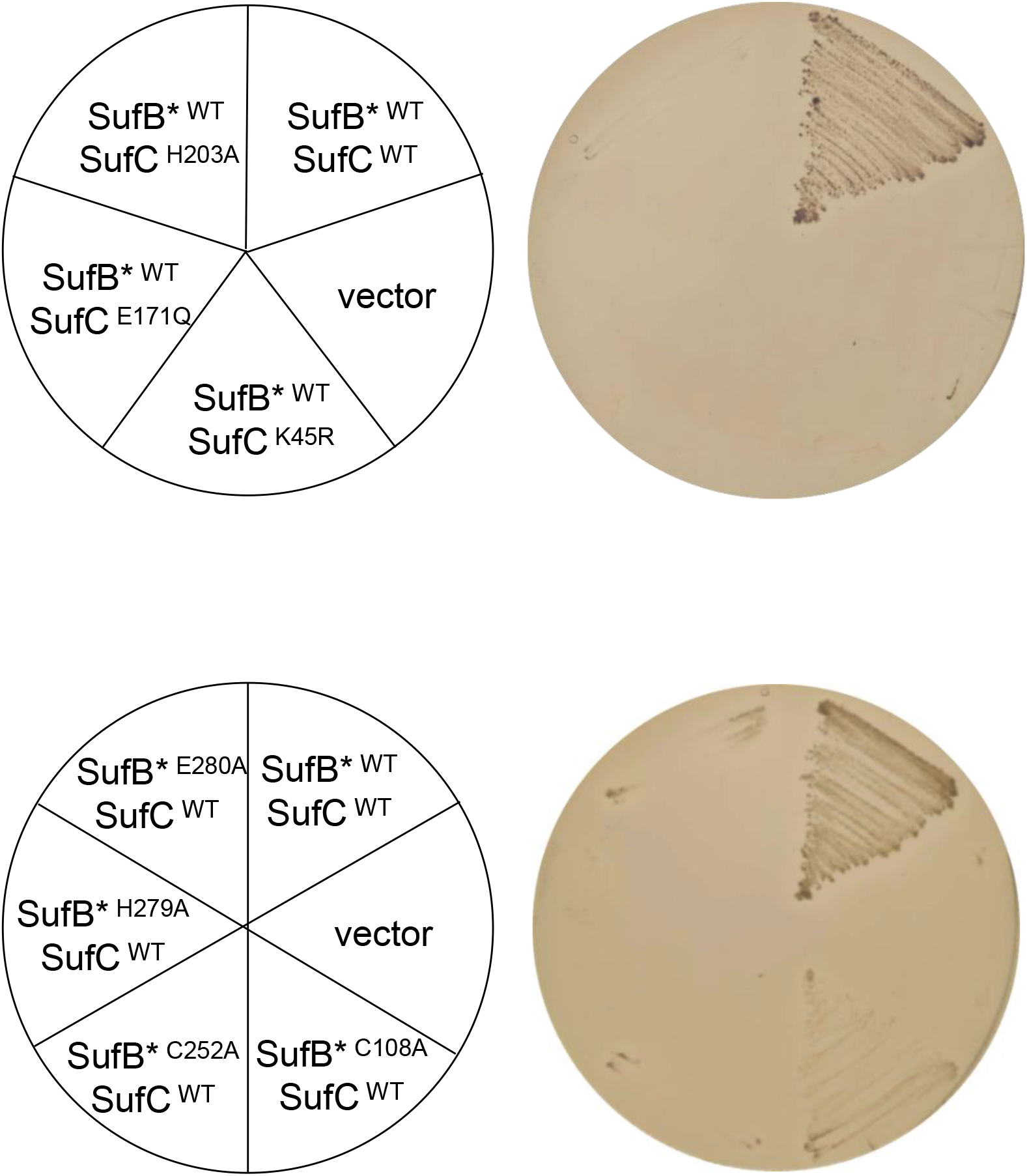
Effect of amino acid substitutions in *M. jannaschii* SufB* or SufC on the heterologous complementation. The UT109 cells harboring the plasmid pUMV22 Sp^r^ were transformed with pBBR-*Mj sufB*C* carrying a point mutation in *sufC* (K45R, E171Q, or H203A) or *sufB\** (C108A, C252A, H279A, or E280A). Cells were grown on LB-glucose plates supplemented with 5 mM Na_2_S (without MVA) at 37°C for 48 hr under anaerobic conditions.

By contrast, amino acid substitution of *Mj* SufB* C108A had only a modest effect; a slow growth phenotype was observed for the complemented cells (Figure 6). The corresponding Cys254 residue in *Ec* SufB plays an essential role in accepting persulfide sulfur from *Ec* SufE (Yuda *et al*., 2017), whereas the nonessential role of *Mj* SufB* Cys108 is consistent with the observation that *Mj* SufB*_2_C_2_ utilizes sulfide as described above. We further examined other amino acids conserved in the SufB* subfamily. However, the alanine substitution of Cys78, Gln88, Asn165, or His249 in *Mj* SufB* did not cause noticeable defects, and the substitution of His118 or His138 caused a slightly retarded growth phenotype in which the complementation was partial (data not shown). Hence, Cys252, His279, and Glu280 are the three essential residues in *Mj* SufB* so far identified. The molecular basis for the selective utilization of sulfide is currently unknown.

### Artificial conversion of *Mj* SufB*_2_C_2_ to an asymmetric SufBC_2_D type complex

In contrast to the asymmetric arrangement of the three functional residues at the *Ec* SufB-SufD interface (Cys405 and Glu434 in *Ec* SufB, and His360 in *Ec* SufD), the corresponding three functional residues in *Mj* SufB* (Cys252, His279, and Glu280) are presumed to be doubled and symmetrically arranged at the interface of the *Mj* SufB* homodimer. In order to investigate whether the symmetric arrangement of the six residues is required for the function of *Mj* SufB*_2_C_2_ or, like *Ec* SufBC_2_D, three asymmetric residues alone are enough, we prepared two copies of *Mj sufB\**, in which C252A and E280A substitutions were introduced into one copy mimicking *Ec* SufD (termed *Mj sufB*1*, which retained the essential His279), and a H279A substitution was introduced into the other, mimicking *Ec* SufB (termed *Mj sufB*2*,which retained the essential Cys252 and Glu280). It should be noted that the *Ec* SufB His433 conserved at the corresponding position (Figure S1d) is not essential and can be functionally substituted with Ala (Yuda *et al*., 2017). Coexpression of the *Mj sufB*1* and *sufB*2* genes with *Mj sufC* from the two compatible plasmids (pRK-*Mj sufB*1* and pBBR-*Mj sufB*2-sufC*) in the *E. coli* UT109 cells resulted in no complementation. However, several spontaneous revertants were generated during the prolonged incubation that restored growth in the absence of MVA under anaerobic and Na_2_S-supplemented conditions. They showed a slightly slower growth compared to the cells harboring wild-type *Mj sufB*C* suggesting partial restoration of Fe-S cluster biosynthesis (Figure 7). Two plasmids were separately recovered from each revertant and subjected to sequence analysis, which revealed one additional point mutation each in the coding region of *Mj sufB*1* or *Mj sufB*2* that caused amino acid substitutions G225S, G284S, A279V, V262F, S264I, or V268M (Figure S5). Homologous recombination between the *Mj sufB*1* and *Mj sufB*2* was not observed at all, which is because the codons for H279 and E280 are next to each other in the *Mj sufB\** sequence. Reintroduction of the plasmids into *E. coli* UT109 cells confirmed that both plasmids were required for the complementation (data not shown). For instance, pBBR-*Mj sufB*2-sufC* harboring the original H279A mutation and an additional G284S mutation in *sufB\** did not complement UT109 unless *Mj sufB*2* harboring the C252A and E280A substitutions was coexpressed, suggesting functional interaction between *Mj* SufB*2 (H279A+G284S) and *Mj* SufB*1 (C252A+E280A). Thus, an additional single amino acid substitution allowed *Mj* SufB*1 and *Mj* SufB*2 to work asymmetrically as an Fe-S cluster biosynthesis scaffold.

**Figure 7.**
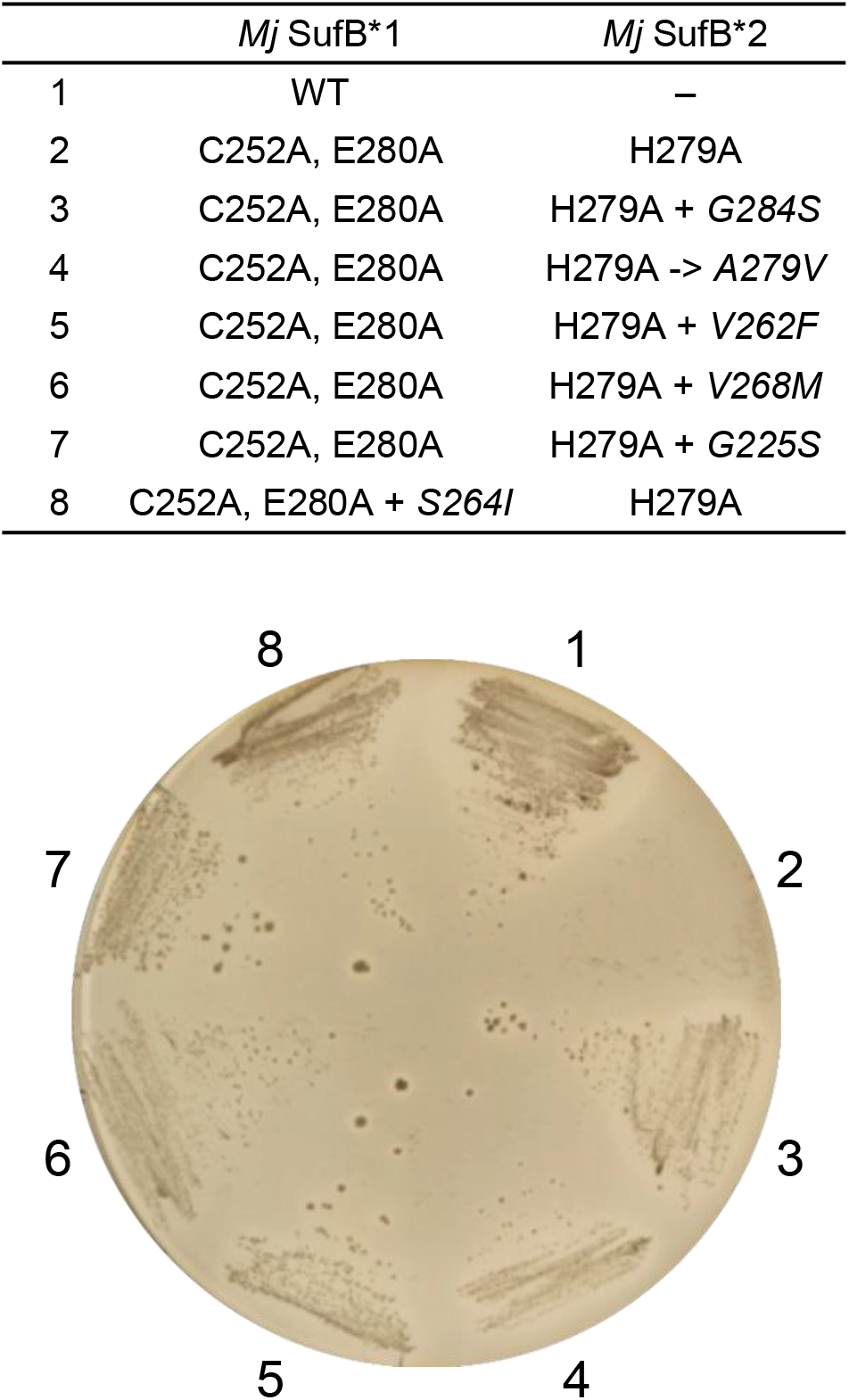
Artificial gene duplication of *M. jannaschii sufB\** and conversion of the symmetric SufB*_2_C_2_ to the asymmetric SufBC_2_D type. The *Mj sufB*1* (carrying the C252A and E280A substitutions) was coexpressed with *sufB*2* (carrying the H279A substitution) and *sufC* from two compatible plasmids in the *E. coli* UT109 cells. Spontaneous revertants originating from these cells carried one additional point mutation each in the coding region of *Mj sufB*1* or *Mj sufB*2* that caused amino acid substitutions G225S, G284S, A279V, V262F, S264I, or V268M (shown in italics). They were grown on LB-glucose plates supplemented with 5 mM Na_2_S (without MVA) at 37°C for 48 hr under anaerobic conditions.

### Biosynthesis systems in Methanothrix thermoacetophila and Archaeoglobus fulgidus

Phylogenetic analysis of the SufB* sequences revealed that they are divided into two deeply branched two groups termed Groups 1 and 2 (Figure S6), where the functionally essential residues in *Mj* SufB* (Cys252, His279, and Glu280) are strictly conserved in both groups (Figure S1d). Despite the tremendous variation in the N-terminal length, the Group 2 sequences including *Mj* SufB* show a defining characteristic of short C-termini, while the Group 1 sequences retain the C-termini corresponding to the last α-helix in *Ec* SufB and SufD. Next, we examined whether the Group 1 *sufB*C* can also complement the *E. coli* mutant UT109 using the synthetic operons cloned from *M. thermoacetophila* (*Mt*) and *A. fulgidus* (*Af*). As in the case of *Mj* SufB*C, the complementation was observed for cells harboring the plasmid pBBR-*Mt sufB*C* or pBBR-*Af sufCB\** under anaerobic conditions only when Na_2_S was supplemented to the media (Figure 8). Partial complementation by *Mt sufB*C* might be due to the rare codons that were not optimized for expression in *E. coli*. Thus, both groups of SufB*C can serve as Fe-S cluster biosynthesis systems using sulfide. We also confirmed that alanine substitution of Cys313, His341 or Glu342 in *Mt* SufB* (corresponding to Cys252, His279 or Glu280 in *Mj* SufB*) also abolished the heterologous complementation (data not shown).

**Figure 8.**
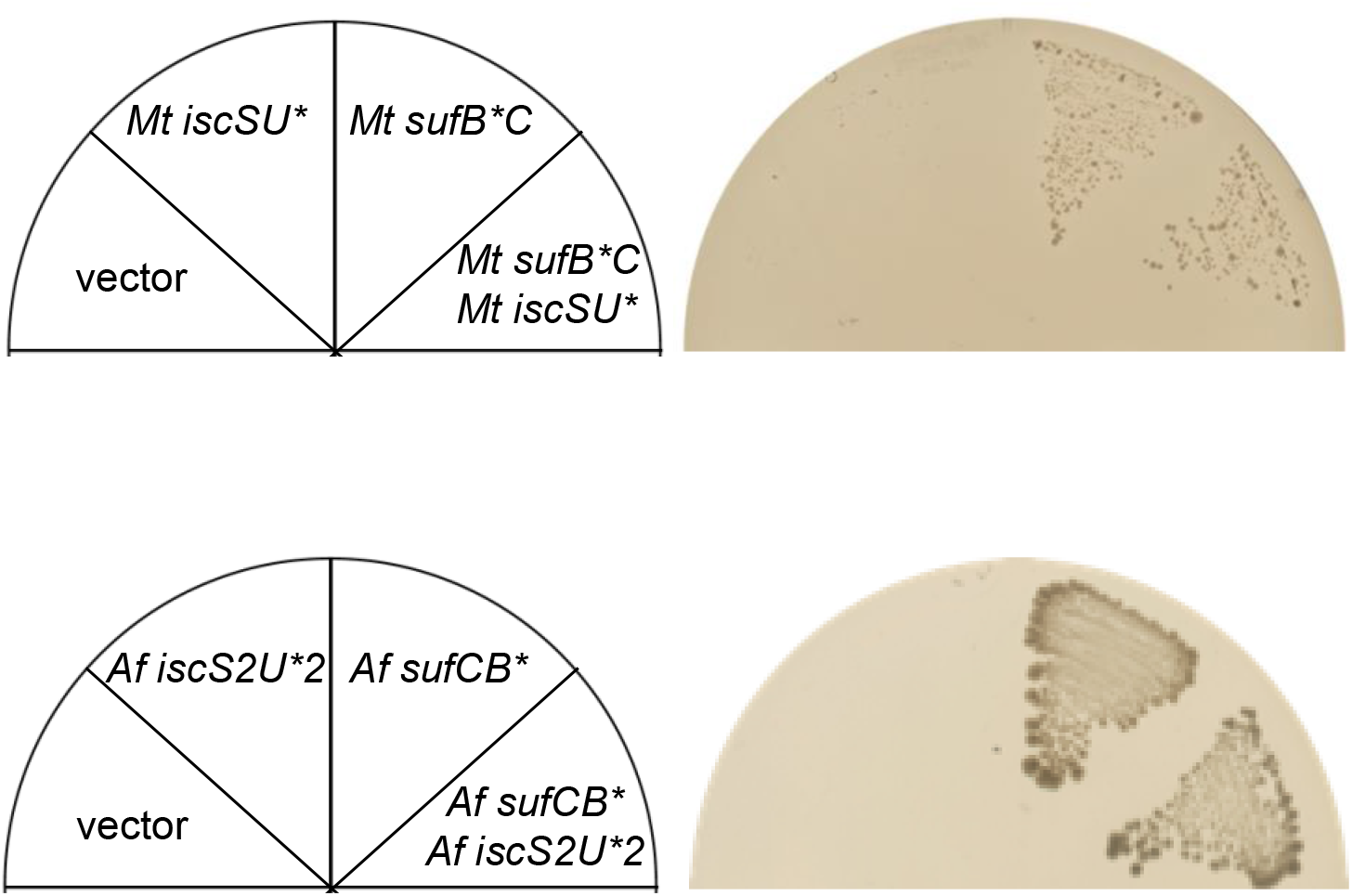
Complementation tests of *E. coli* UT109 by the Group 1 *sufB*C* and/or minimal *iscSU\** from *M. thermoacetophila* (*Mt*) and *A. fulgidus* (*Af*). The UT109 cells harboring pUMV22 Sp^r^ were transformed with plasmids carrying *Mt sufB*C* or *Af sufCB\** and/or plasmids carrying *Mt iscSU\** or *Af iscS2U*2.* Cells were grown on LB-glucose plates supplemented with 5 mM Na_2_S (without MVA) at 37°C for 72 hr under anaerobic conditions.

More than half of the organisms that harbor the *sufB*C* genes (including *M. thermoacetophila* and *A. fulgidus*) also harbor the *iscS* and *iscU\** genes for the minimal ISC system (Figures 1 and S6). Although it has been speculated that there is sulfur transfer interaction between IscSU* and SufB*C, amino acid sequences of IscU* are clearly distinguished from that of SufU (sulfur-transfer protein) in that amino acid residues critical for *de novo* Fe-S cluster assembly in *E. coli* IscU (Cys37, Asp39, Cys63, Lys103, His105 and Cys106) are strictly conserved (Figure S1b). Likewise, the IscS sequences belong to the Type I cysteine desulfurase group (Figure S1a), but *Af* IscS1 and IscS2 are exceptional in that the proteins have no cysteine desulfurase activity due to the substitution of the PLP-binding Lys residue to Asp (at the position of 199 in *Af* IscS2). It has been proposed that *Af* IscS functions solely as a scaffold protein, providing one Cys ligand to bind to the [2Fe-2S] cluster onto the *Af* IscS-IscU complex, but does not provide sulfur (Pagnier *et al*., 2015), where the amino acid sequences are 85% identical between *Af* IscS1 and IscS2, and 100% between *Af* IscU*1 and IscU*2. We constructed synthetic operons for *Mt iscSU\** and *Af iscS2U*2*, but neither procured functional complementation of the *E. coli* mutant UT109 even when anaerobic conditions were combined with Na_2_S supplementation (Figure 8). We also examined the coexpression of *Mt sufB*C* and *Mt iscSU\** from two plasmids, but no stimulatory effect was observed compared to *Mt sufB*C* alone. Without the Na_2_S supplementation, no complementation was observed even when *Mt sufB*C* and *Mt iscSU\** were coexpressed (data not shown). Similarly, the coexpression of *Af sufCB\** and *Af iscS2U*2* from two plasmids did not improve the heterologous complementation by *Af sufCB\** alone. Therefore, our attempts to experimentally clarify the *in vivo* function of IscSU* have been unsuccessful.

## Discussion

### SufB*C serves as an *in vivo* Fe-S cluster biosynthesis machinery using inorganic sulfide

Some bacteria and archaea encode only two components, SufB* and SufC, but lack the genes for SufD and the sulfur supply system (SufSE or SufSU) that usually support Fe-S cluster biosynthesis. In this study, we report for the first time that SufB*C from *M. jannaschii*, *M. thermoacetophila* and *A. fulgidus* can serve as an Fe-S cluster biosynthesis system *in vivo*. In heterologous complementation experiments using an *E. coli* mutant, the archaeal SufB*C systems were able to substitute for the well-characterized *Ec* SUF and ISC systems, where the recovery of a MVA-independent growth phenotype demonstrated the restoration of the activity of Fe-S enzymes IspG and IspH, which are involved in the isoprenoid biosynthesis MEP pathway. Restoration of the activity of another Fe-S enzyme, FDH_H_, was also demonstrated for the cells complemented by *Mj* SufB*C. Most importantly, the restoration was observed only when sulfide was supplemented to the medium under anaerobic conditions (Figure 4). This is consistent with the habitat of *M. jannaschii*, *M. thermoacetophila* and *A. fulgidus*, which are obligate anaerobes growing in environments rich in sulfide, where *A. fulgidus* also reduces sulfate to sulfide (Klenk *et al*., 1997). In another anaerobic archaeon, *M. maripaludis*, that contains *sufCB\** as essential genes, isotope labeling experiments have suggested that this methanogen uses exogenous sulfide as the sulfur source for Fe-S clusters instead of L-cysteine (Liu *et al*., 2010). At neutral pH, one-third of sulfide is in nonionized form (H_2_S) and may freely diffuse across cell membranes (Liu *et al*., 2012). It is currently unknown whether the H_2_S or HS^-^ inside the cells is directly utilized by SufB*C or not. In view of the high affinity between HS^-^ and ferrous iron, it is tempting to speculate that they first associate in the cytosol to form aqueous iron-sulfide complexes (Rickard & Luther, 2007), which are then assembled into biologically available Fe-S clusters by SufB*C. Recent studies on *Methanococcus voltae* and *Methanosarcina barkeri* (both harboring the *sufCB\** and *iscSU\** genes) have demonstrated that these methanogens can reductively dissolve extracellular Fe_2_S (pyrite) and utilize the dissolution products to meet Fe and S nutritional demands (Payne *et al*., 2021; Spietz *et al*., 2022). In marked contrast to SufB*C, the canonical Fe-S cluster biosynthesis systems (NIF, ISC, and SUF) so far characterized utilize L-cysteine as the sulfur source for the Fe-S cluster. In *E. coli*, the sulfur supply system (SufS and SufE) was absolutely required for the SUF machinery even when sulfide was supplemented to the medium under anaerobic conditions (Figure S3). Thus, the major difference between SufB*C and SufBCD is that the former utilizes inorganic sulfide and the latter the sulfur derived from L-cysteine, although both systems appear to serve as a core scaffold for the *de novo* assembly of Fe-S clusters.

AlphaFold2 predicts a structural model of SufB*_2_C_2_ not only for *Mj* SufB*C but also for the complexes from *M. thermoacetophila* and *A. fulgidus* (Figure 5). Overall, the architecture of SufB*_2_C_2_ is similar to that of the *Ec* SufBC_2_D complex. According to the mechanistic model of the *E. coli* SUF machinery, the sulfur derived from L-cysteine is transferred from SufS to SufE in the form of persulfide (-SSH) and then to SufB Cys254 in the SufBC_2_D complex (Layer *et al*., 2007; Yuda *et al*., 2017). Most probably, the Fe-S cluster is assembled at the interface of SufB and SufD using three essential residues (Cys405 and Glu434 in *Ec* SufB and His360 in SufD). The conformational change of this region is a prerequisite for the three residues to work in concert, and this is probably induced by ATP driven dimerization of two subunits of SufC (Hirabayashi *et al*., 2015). The functional residues of the *Ec* SufBC_2_D complex are conserved in almost all SufB*C sequences, and in this study, the essential role of Lys45 (Walker A motif), Glu171 (Walker B motif), and His203 (H-motif) in *Mj* SufC and Cys252, His279, and Glu280 in *Mj* SufB* was confirmed by site-directed mutations (Figure 6). One exception is the *Mj* SufB* Cys108 corresponding to the *Ec* SufB Cys254 that receives persulfide sulfur from *Ec* SufE, but this is not surprising considering that *Mj* SufB*C utilize inorganic sulfide instead of L-cysteine. The scaffold mechanism for the Fe-S cluster assembly appears to be conserved between SufB*_2_C_2_ and SufBC_2_D although they use different sulfur sources.

Although the SufB_2_C_2_ and the SufC_2_D_2_ complexes have been observed in *E. coli* in the absence of SufD and SufB, respectively (Saini *et al*., 2010; Wada *et al*., 2009), they did not show any competence for *in vivo* Fe-S cluster biosynthesis (Figure S3). In contrast to the asymmetric SufB-SufD interface of the functional SufBC_2_D complex, SufB*_2_C_2_ appears to adopt a symmetric SufB*-SufB* interface where three functional residues (Cys252, His279, and Glu280 in *Mj* SufB*) are doubled. Attempts to modify *Mj* SufB*_2_C_2_ using two copies of *Mj sufB*\* to resemble the asymmetric SufBC_2_D were not successful per se, namely, *Mj* SufB*1 carrying C252A and E280A substitutions (mimicking *Ec* SufD) and *Mj* SufB*2 carrying H279A substitution (mimicking *Ec* SufB) do not function independently of each other, nor do they function in combination (in the presence of *Mj* SufC). However, an additional suppressor mutation allowed the two variant *Mj* SufB* molecules to work in concert and achieve heterologous complementation (Figure 7). Intriguingly, the amino acid substitutions (G225S, G284S, A279V, V262F, S264I, or V268M) responsible for the suppressor function are likely lined up at the interface of the *Mj* SufB*1-SufB*2 in the proximity of the three essential residues Cys252, His278, and Glu280 (Figure S5). It is also noteworthy that all these substitutions resulted in bulkier side chains. Although the precise role of these suppressor mutations is currently unknown, we speculate that the substitutions may facilitate the conformational change of the SufB*1-SufB*2 interface that is proposed for *Ec* SufB-SufD (Hirabayashi *et al*., 2015). Thus, it was experimentally possible, by introducing just a few amino acid substitutions, to convert the symmetric SufB*_2_C_2_ structure to the asymmetric SufBC_2_D type.

### Evolutionary trajectory of the SUF system

Genes for SufB*C are distributed in both archaea and bacteria, and according to our literature search, all of them are anaerobes. They are mainly found in the Euryarchaeota including methanogens and in the bacterial classes of Clostridia and Deltaproteobacteria, but also turn up in the archaeal TACK group as well as in a tremendous variety of bacterial species (Figure S6). The Group 1 SufB* sequences with long C-termini are common in archaea, whereas the Group 2 sequences (short C-termini) occur in bacteria, although there is no clean split between the two domains. In the present study, we have demonstrated characteristics common to the Group 1 and 2 SufB*C systems; both can serve as the Fe-S cluster biosynthesis machinery in *E. coli* cells using exogenously supplemented sulfide as the sulfur source. It has previously been suggested that the ancestral SufB*C system was present in the last universal common ancestor (LUCA) in a low oxygen primordial biosphere and has evolved through repeated horizontal gene transfers among the anaerobic archaea and bacteria (Garcia *et al*., 2022). Even now, this system is vulnerable to oxygen, which may be due to the intrinsic properties of the Fe-S clusters (Imlay, 2006; Lu & Imlay, 2021), and also because sulfide can be directly oxygenated, restricting its distribution to anoxic organisms.

Duplication of SufB* and subsequent mutations resulted in the asymmetric SufBC_2_D system. It is not clear whether this happened in the archaeal or bacterial lineage, since the SufBC_2_D system is also distributed in both domains as is the SufB*C system. In contrast to the SufB*C system, which is distributed only among anaerobes, the SufBC_2_D system is also found among aerobes and facultative anaerobes. These observations suggest that the conversion from the symmetric SufB*_2_C_2_ to the asymmetric SufBC_2_D conveys an advantage in adaptation to the oxygenic environments. In this study, however, the *E. coli* cells complemented by the asymmetric *Mj sufB1*, *sufB2* (carrying the suppressor mutation) and *sufC* were still sensitive to oxygen (data not shown). Outten and coworkers have postulated that the emergence of the asymmetric SufBC_2_D might be related to the bioavailability of iron that is restricted in aerobic habitats (Boyd *et al*., 2014), but this hypothesis also needs to be examined experimentally.

In contrast to the universal employment of the sulfur-supply system (SufSU or SufSE) in the bacterial SUF machinery, the archaeal genomes only scarcely encode the homologs of Type II cysteine desulfurase (SufS) and the sulfur-transfer protein (SufU or SufE). For instance, the facultative anaerobic archaeon *Thermoplasma acidophilum* possesses the genes for *sufCBD* but lacks the sulfur supply system (SufSE and SufSU, Figure 1), leaving open what might be the source of sulfur for the archaeal SufBC_2_D system, in particular in the species thriving under aerobic, non-sulfide-rich environments (Iwasaki, 2010). In the bacterial lineage harboring *sufBCD*, SufSU is much more widely distributed than SufSE, suggesting that SufSU may have been adopted first, and then, SufU was replaced in its role by SufE in some bacterial lineage including cyanobacteria and alpha- and gamma-proteobacteria during the evolutionary history of the SUF system (Figure 2b) (Yokoyama *et al*., 2018). In support of this view, we note that *E. coli* SufSE is more resilient than *Bacillus subtilis* SufSU in the presence of reactive oxygen species such as H_2_O_2_ (our unpublished results).

The minimal ISC system (IscSU*) is also distributed in anaerobes including archaea and bacteria, and it has been suggested that it represents as the prototype of the ISC machinery (Sato *et al*., 2021; Yokoyama *et al*., 2018). Auxiliary components (Fdx, HscA, HscB, and IscA) may then have been recruited to IscSU* during evolution in a process of adaptation to oxygenic environments (Figure 2a). Whereas only certain anaerobic bacteria (mainly Clostridia) possess no more than the IscSU* system, co-occurrence of IscSU* and SufB*C is seen in both anaerobic archaea and bacteria. For instance, the methanogenic archaeon *M. acetivorans* contains two copies of *sufCB\** and three copies of *iscSU\**, among which a redundant role for Fe-S cluster biosynthesis has been proposed from genetic studies (Deere *et al*., 2020; Saini *et al*., 2023). In our *in vivo* experiments, however, no complementation of the *E. coli* mutant was observed by *iscSU\** from *M. thermoacetophila* and *A. fulgidus*. Furthermore, coexpression of *iscSU\** and *sufB*C* showed almost no effect on the complementation compared to *sufB*C* alone. Thus, further studies are needed to find out what is missing from the experiments and to understand how IscSU* can function, apparently without the other known auxiliary components of the canonical ISC machinery. Another important question remains unsolved: what is the cause of the difference between the *in vivo* biosynthesis and the *in vitro* chemical reconstitution of the Fe-S clusters? Living organisms require a dedicated system for the assembly of the Fe-S clusters even in anaerobic and sulfide-rich environments, whereas *in vitro* chemical reconstitution can occur spontaneously as long as ferrous ion and thiols such as glutathione are supplied. In anaerobic organisms like *M. jannaschii*, which thrive around hydrothermal vents enriched in sulfide and ferrous iron, it is unlikely that iron deficiency is the major cause of the requirement for SufB*C, because in such an anoxic environment, iron-induced cytotoxicity would be low. This fundamental question needs to be addressed in future research along with the reaction type and mechanism by which the SufB*_2_C_2_ complex operates.

### Experimental procedures

#### Bacterial strains and cell growth

The *E. coli* strains and plasmids used in this study are listed in Table 1. The mutant strain UT109 is a derivative of *E. coli* MG1655 in which the chromosomal *iscUA-hscBA* and *sufABCDSE* are deleted and which, in the presence of the pUMV22 Sp^r^ plasmid, grows slowly on LB agar plate supplemented with 0.4% glucose and 0.3 mM MVA (mevalonolactone, Sigma-Aldrich) with an absolute dependence on MVA (Tanaka *et al*., 2016). For liquid growth, the cells were cultivated in Superbroth (3.2% bacto tryptone, 2% yeast extract and 0.5% NaCl) supplemented with glucose and MVA. When required, ampicillin (Ap), tetracycline (Tc) and spectinomycin (Sp) were added at concentrations of 50, 5 and 40 µg/ml, respectively. For anaerobiosis, cells were grown on LB agar plates containing 0.4% glucose using the AnaeroPack-Anaero System (Mitsubishi Gas Chemical). Where indicated, Na_2_S was added at a concentration of 2∼10 mM to the melted agar media after autoclaving and cooling to about 50°C. Upon solidification, the plates were used within 30 min. The FDH_H_ activity was assessed by the benzyl viologen agar overlay method (Mihara *et al*., 2002).

**Table 1.** *E. coli* strains and plasmids used in this study.

| Strain | Genotype | Source |
| --- | --- | --- |
| MG1655 | Wild type | Laboratory strain |
| UT109 | MG1655 $\Delta(iscUA-hscBA)::Km^r \Delta(sufABCDSE)::Gm^r$ | (Tokumoto *et al.*, 2004) |
| Plasmid | Description | Source |
| pUMV22 Sp<sup>r</sup> | Sp<sup>r</sup>; pUC118 derivative carrying three genes for MVA kinase, PMVA kinase, and DPMVA decarboxylase | (Tanaka *et al.*, 2016) |
| pBBR1MCS-4 | Ap<sup>r</sup>; pBBR replicon, low-copy-number vector, P<sub>lac</sub> promoter | (Kovach *et al.*, 1995) |
| pBBR1MCS-3 | Tc<sup>r</sup>; pBBR replicon, low-copy-number vector, P<sub>lac</sub> promoter | (Kovach *et al.*, 1995) |
| pBBR- *Mj sufB* * | pBBR1MCS-4 derivative carrying *M. jannaschii sufB* * | This study |
| pBBR-*Mj sufC* | pBBR1MCS-4 derivative carrying *M. jannaschii sufC* | This study |
| pBBR- *Mj sufB* *C | pBBR1MCS-4 derivative carrying *M. jannaschii sufB* *C | This study |
| pBBR- *Mj sufB* *2C | pBBR- *Mj sufB* *C derivative harboring H279A substitution | This study |
| pRKNMC | Tc<sup>r</sup>; IncP-1 replicon, low-copy-number vector, P<sub>lac</sub> promoter | (Nakamura *et al.*, 1999) |
| pRK- *Mj sufB* *1 | pRKNMC derivative carrying *M. jannaschii sufB* * harboring C252A and E280A substitutions | This study |
| pRK-*Mj apbC* | pRKNMC derivative carrying *M. jannaschii apbC* | This study |
| pBBR- *Mt sufB* *C | pBBR1MCS-3 derivative carrying *M. thermoacetophila sufB* *C | This study |
| pBBR- *Mt iscSU* * | pBBR1MCS-4 derivative carrying *M. thermoacetophila iscSU* * | This study |
| pBBR- *Af sufCB* * | pBBR1MCS-4 derivative carrying *A. fulgidus sufCB* * | This study |
| pRK- *Af iscS2U* *2 | pRKNMC derivative carrying *A. fulgidus Af iscS2U* *2 | This study |
| pRKSUF017 | pRKNMC derivative carrying *E. coli sufABCDSE* | (Takahashi & Tokumoto, 2002) |
| pRK-*Ec sufA_CDSE* | pRKSUF017 derivative carrying *E. coli sufACDSE* ($\Delta sufB$) | (Yokoyama *et al.*, 2018) |
| pRK-*Ec sufAB_DSE* | pRKSUF017 derivative carrying *E. coli sufABDSE* ($\Delta sufC$) | (Hirabayashi *et al.*, 2015) |
| pRK-*Ec sufABC_SE* | pRKSUF017 derivative carrying *E. coli sufABCSE* ($\Delta sufD$) | (Wada *et al.*, 2009) |
| pRK-*Ec sufABCD_E* | pRKSUF017 derivative carrying *E. coli sufABCDE* ($\Delta sufS$) | (Yokoyama *et al.*, 2018) |
| pRK-*Ec sufABCDS* | pRKSUF017 derivative carrying *E. coli sufABCDS* ($\Delta sufE$) | (Takahashi & Tokumoto, 2002) |
| pRK-*Ec sufABCD* | pRKSUF017 derivative carrying *E. coli sufABCD* ($\Delta sufSE$) | (Yokoyama *et al.*, 2018) |

#### Plasmid construction

Sequences of PCR primers are listed in Table S1. The nucleotide sequences of the synthetic genes and operons are also shown in “Supporting Information”. These genes were cloned in the plasmids and expressed in *E. coli* under the control of the lactose promoter.

The coding regions for *M. jannaschii sufB\** and *sufC* were synthesized so as to optimize the codons for expression in *E. coli* cells (GenScript). The *Mj sufC* fragment was digested by NdeI and BamHI and ligated into the corresponding sites of pET-21a (Novagen-Merck Millipore). The *Mj sufC* coding region (without His_6_ tag sequence) with the upstream ribosome binding sequence was excised by XbaI and SacI digestion and ligated into the corresponding sites of pBBR1MCS-4 (Kovach *et al*., 1995). The synthesized *Mj sufB\** fragment was digested by BamHI and HindIII and ligated into the corresponding sites of pACYCDuet-1 (Novagen-Merck Millipore). The coding region of *Mj sufB\** with the N-terminally fused His_6_ sequence and the upstream ribosome binding sequence were amplified by PCR using the primers SalI-His-MjSufB-for and HindIII-His-MjSufB-rev. The amplified fragment was digested by SalI and HindIII and cloned into the corresponding sites of pBBR-*Mj sufC* yielding the pBBR-*Mj sufB*C.* The *Mj sufB\** fragment was excised by KpnI and EcoRI digestion and subcloned into the corresponding sites of pRKNMC (Nakamura *et al*., 1999) yielding the pRK-*Mj sufB*.* Site-directed mutations were introduced into the pBBR-*Mj sufB*C* or pRK-*Mj sufB\** plasmids by inverse PCR using the appropriate mutagenic primers.

For the construction of pRK-*Mj apbC*, the coding region of *apbC* was amplified by PCR using the genomic DNA of *M. jannaschii* as a template and the primers Mj-apbC-Fw-Nd and Mj-apbC-Rv-Xh. The amplified fragment was digested by NdeI and XhoI and ligated into the corresponding sites of pRSFDuet-1 (Novagen-Merck Millipore). The coding region of *Mj apbC* and the upstream ribosome binding sequence were amplified by a second PCR using the primers Mj-apbC-Fw2-Sc and Mj-apbC-Rv-Xh. The amplified fragment was digested by SacI and XhoI and cloned into the corresponding sites of pRKNMC.

For the construction of pBBR-*Mt sufB*C*, the coding region of *Mt sufB\** with the artificial ribosome binding sequence was amplified by PCR using the genomic DNA of *M. thermoacetophila* as a template and the primers Mt-sufB*-F-Sl and Mt-sufB*-R-Hd. The amplified fragment was digested by SalI and HindIII and ligated into the corresponding sites of pBBR1MCS-4. The coding region of *Mt sufC* with the artificial ribosome binding sequence was amplified by PCR using the primers Mt-sufC-F-Hd and Mt-sufC-R-EcXb. The amplified fragment was digested by HindIII and XbaI and cloned into the corresponding sites of pBBR-*Mt sufB\**. The SalI-XbaI fragment harboring *Mt sufB*C* was excised and subcloned into pBBR1MCS-3.

For the construction of pBBR4-*Mt iscSU\**, the coding region of *Mt iscS* with the artificial ribosome binding sequence was amplified by PCR using the genomic DNA of *M. thermoacetophila* as a template and the primers XhoI-SD-Mt IscS-fw and Mt IscS-stop-HindIII-rv. The amplified fragment was digested by XhoI and HindIII and ligated into the corresponding sites of pBBR1MCS-4. The coding region of *Mt iscU\** (without His_6_ tag sequence) with the upstream ribosome binding sequence was excised from pET-21a-*Mt iscU\** (Kunichika *et al*., 2021) by XbaI and SacI digestion and cloned into the corresponding sites of pBBR1-*Mt iscS*.

The operons for *A. fulgidus sufCB\** and *iscS2U*2* were synthesized so as to optimize the codons for expression in *E. coli* cells with artificial ribosome binding sequences (Twist Bioscience). The HindIII-SacI fragment harboring *Af sufCB\** was subcloned into pBBR1MCS-4 and that of *Af iscS2U*2* into pRKNMC.

#### Expression and purification of SufB*C proteins

*M. jannaschii* SufB*C proteins (wild type and mutated variants) were expressed from the pBBR-*Mj sufB*C* plasmid in *E. coli* MG1655 cells. The cells were cultivated in Terrific Broth for 20 hr at 20°C under induction by 1.0 mM isopropyl-β-D-thiogalactopyranoside. Harvested cells were disrupted by sonication. After centrifugation, the supernatant fraction was loaded onto a HisTrap^TM^ FF column (GE Healthcare) and washed with the buffer containing 50 mM Tris-HCl (pH7.8), 0.5 M KCl, 1 mM DTT, and 20 mM imidazole. The proteins were eluted with the washing buffer containing 250 mM imidazole. The purified *Mj* SufB*C and the variant forms exhibited no signs of Fe-S clusters.

## Acknowledgements

This work was supported by JSPS KAKENHI Grant Numbers JP20H03204 (to Y.T.), JP21K19204 (to Y.T.), JP17K14510 (to T.F.), and JP20H03196 (to K.W.).

## Data Availability Statement

The data that support the findings of this study are available from the corresponding author upon reasonable request.

## Author contributions

T.F., E.Y., K.K., and Y.T. designed research; M.M., T.M., N.M., X.Z., N.K., S.M., K.K., and S.O. performed research; M.M., T.M., S.O., T.F., K.W., and Y.T. analyzed data; K.W., T.F., and Y.T. wrote the paper.

