## Supporting Information for "The minimal SUF system can substitute for the canonical iron-sulfur cluster biosynthesis systems by using inorganic sulfide as the sulfur source"

**Table S1. Oligonucleotides used in this study.**

| Oligonucleotide | Sequence |
|---|---|
| SalI-His-Mj SufB*-for | TTTTGTCGACCTTTAATAAGGAGATATACCATGGGCAGC |
| HindIII-His-Mj SufB*-rev | CCGCAAGCTTTTACAGATCACCAATCA |
| C108A-Mj SufB*-for | GCTTCATTTCCGAAAGGCAAAGGTATCAAAC |
| C108A-Mj SufB*-rev | ATGGGACATCAGACTAATAGAGCTGT |
| C252A-Mj SufB*-for | GCTGCAGAAATCGTGAAAGGCAACGC |
| C252A-Mj SufB*-rev | GTCAATATGACCTTTAGAATACGGCG |
| H279A-Mj SufB*-for | GCAGAAGCAGCTATCGGTTCAGTTGATAA |
| H279A-Mj SufB*-rev | GGTAATGCGTGCTTTGTCATCACG |
| E280A-Mj SufB*-for | GCAGCAGCTATCGGTTCAGTTGATAAAAAAC |
| E280A-Mj SufB*-rev | GTGGGTAATGCGTGCTTTGTCATC |
| K45R-Mj SufC-for | AGAAGTACCCTGGCCTATACGATTATG |
| K45R-Mj SufC-rev | ACCCGCGCCATTCGGGCCGATA |
| E171Q-Mj SufC-for | CAACCGGACAGCGGCATCGATATT |
| E171Q-Mj SufC-rev | ATCCAGGATGGCCAGATCCGG |
| H203A-Mj SufC-for | GCTCGTGAAGAACTGGCGGAACAC |
| H203A-Mj SufC-rev | CGTGATGACCAGCAGGGAGCAA |
| Mj-apbC-Fw-Nd | GCCGCGCATATGGCTGAGTGTGATGGAAAA |
| Mj-apbC-Rv-Xh | GGCCCTCGAGTCATTCTTTTTTACCTTCGACCTTT |
| Mj-apbC-Fw2-Sc | GCCGAGCTCAGCGGATAACAATTCCCCAT |
| Mt-sufB*-F-Sl | GCGCGTCGACAAGGAGATATACATATGTCCGACATAGCTGAGAG |
| Mt-sufB*-R-Hd | GCGCAAGCTTCCCGCCCTCGTTTTAGGC |
| Mt-sufC-F-Hd | GCGCAAGCTTAAGGAGACACACTAATGTTCGGGCTGCTCGAGAT |
| Mt-sufC-R-EcXb | GCGCTCTAGAATTCTATGTCGGACATAGGCACC |
| XhoI-SD-Mt IscS-fw | CCCGCTCGAGAAGGAGATATACCATGAGGCGCATATACATGGACC |
| Mt IscS-stop-HindIII-rv | CCCGAAGCTTCTACCTCAGCGCTGATATGCCGCGGAAC |

**Nucleotide sequences of the synthetic genes and operons expressed in *E. coli*.** *M. jannaschii sufB*C* (a) and *apbC* (b), *M. thermoacetophila sufB*C* (c) and *iscSU** (d), and *A. fulgidus sufCB** (e) and *iscS2U*2* (f). The coding regions are underlined.

(a) *Methanocaldococcus jannaschii sufB*C*

GTCGACCTTTAATAAGGAGATATACCATGGGCAGCAGCCATCACCATCATCACCACAGCCAGGATCCGAGCATTAAAGAAGAACTGAT
GGAAATTATCGAAGCTATCAAATACACCAGCGAAAAACCGGAAGAAATCGTCCACGGCAAAGGTCCGCGTATTATCGTCAAAGAATCG
CGCATTATCGATGTGCAGGGCGACGAAGGTATTATCCTGGAAGGCAAAGAAGAAGATGGTAAAATCAAAGCGAAAATCATCGTGAAGA
AAGGCTACAAATTCAAATACCCGATCCACATGTGCTTCGGTATTACCGAAGAAAACATCAGCCAAATTATCGATGTCGAAATTATCCT
GGAAGAAGACAGCTCTATTAGTCTGATGTCCCATTGTTCATTTCCGAAAGGCAAAGGTATCAAACACATCATGAACGGCATCATCAAA
ATCGGTAAAAACGCCAAATTCTCCTACAACGAATTTCATTACCACGGCATGGATGGTGACATTCTGGTTAAACCGACCGTTAAAGTCG
AAATCGATGAAGGCGGTATTTATATCTCAAACTTCACCCTGACGAAAGGCCGTATTGGCACCCTGGACATCGAACAGGAAATTATCGC
GAAGAAAGATGCCATTATCGACATTACCACGCGCACGTACGCGATCAAAGAAGATGTGGTTAAAGTGAATGAAGTCGTGAAACTGAAC
GGCGAAAATGCCAAATGCATTATCAAATCTCGTGGTGCGGCGATGGATAACTCGAAAATTAGCCTGAAACTGAAAATCGAAGGCAATG
CGCCGTATTCTAAAGGTCATATTGACTGTGCAGAAATCGTGAAAGGCAACGCTGAAGTTGAAAGTATTCCGATTGTGGTCGTGCGTGA
TGACAAAGCACGCATTACCCACGAAGCAGCTATCGGTTCAGTTGATAAAAAACAACTGGAAACGCTGATGGCTAAAGGTCTGGACGAA
GATGAAGCCACGGAAATTATTGTCAAAGGCATGATTGGTGATCTGTAAAAGCTTGATATCGAATTCCTGCAGCCCGGGGGATCCACTA
GTTCTAGAAATAATTTTGTTTAACTTTAAGAAGGAGATATACATATGGTGTCTATTATGCTGCTGAAAGTGGAAGACCTGCATGTTTA
TCGTGGCAACCGTGAAATCCTGAAAGGCGTGAATCTGACCGTCGAAGAAAACGAAATTCATGCAATTATCGGCCCGAATGGCGCGGGT
AAAAGTACCCTGGCCTATACGATTATGGGCATCTCCGGTTACAAACCGACCAAAGGCCGTATCATCTTCAAAGGTGTGGATATTATCG
ACAAAAACATTACCGAACGTGCGCGCATGGGTATGACGCTGGCATGGCAGGAACCGGCTCGCTTCGAAGGCATCAAAGTTAAAAACTA
TCTGATGCTGGGTATGAACGAAAAATACAAGAAAGATAAAGAAATCGCAGAAGAAAAAATCCGTGAAGCTCTGAAACTGGTTAATCTG
GACCCGGACAAATATCTGGATCGCTACGTGGATGAAACCCTGAGCGGCGGTGAACGTAAACGCATTGAACTGGCGTCTATTATCTGCA
TGGAACCGGATCTGGCCATCCTGGATGAACCGGACAGCGGCATCGATATTGTGTCTTTTGACGAAATCAAACGTGTTTTCGATTACCT
GAAAGACAAAGGTTGCTCCCTGCTGGTCATCACGCATCGTGAAGAACTGGCGGAACACGCGGATCGTGTCTCACTGATTTGTGCGGGC
GAAGTGATCAAATCGGGTGATCCGAAAGAAGTTGGTGAATTTTACAAAAAAGAATGCGGCAAATGCTACAAAAAAGTTCCGGACGGCA
AATAACTCGAGATCGGATCCGAATTCGAGCTC

(b) *Methanocaldococcus jannaschii apbC*

GAGCTCAGCGGATAACAATTCCCCATCTTAGTATATTAGTTAAGTATAAGAAGGAGATATACATATGGCTGAGTGTGATGGAAAATGT
GACACTTGTCCATCAAAAAATACCTGCCCAGATACAAAGAAACTCTTAGCCCAGCAAGATGCAAAAATTAGAGAAAATATGTCAAAAA
TAAAACATAAAATAGTTATTTTGAGTGGTAAAGGAGGGGTTGGGAAATCAACAGTAACAGTTAATTTAGCTGCTGCTCTAAATTTAAT
GGGCAAAAAGGTTGGAGTTTTAGATGCGGATATTCACGGCCCTAACATTCCAAAGATGCTTGGGGTTGAGAACACCCAACCTATGGCA
GGACCAGCTGGAATATTTCCAATAGTTACAAAAGATGGAATAAAAACCATGTCTATTGGATATCTATTACCAGATGACAAAACTCCTG
TTATTTGGAGGGGGCCAAAGGTTAGCGGAGCTATTAGGCAATTTCTATCAGATGTAGTTTGGGGAGAACTTGATTATTTATTAATAGA
TACTCCTCCAGGGACAGGAGATGAGCAATTAACTATCATGCAATCAATTCCAGATATTGATGGAGCTATAATTGTAACAACACCAGAA
GAAGTTTCTGTCTTGGATGTTAAAAAATCCATTATGATGGCTAAAATGCTAAACATCCCAATTATTGGAATTATTGAAAATATGAGCG
GGTTTGTTTGCCCATACTGCAATAAAGTTGTGGATATATTTGGTAGAGGAGGGGGAGAAAAAGCTGCTAAAGAGCTTGGAGTTGAATT
TTTAGGTAGAATTCCTTTAGATATTAAAGCAAGAGAAGCAAGTGATAAAGGAATTCCAATGGTTTTACTTGATTGTAAAGCAAGTGAA
GAGTTTAAAAAGATTGTTAAAAGAATTGTTGAAAAGGTCGAAGGTAAAAAAGAATGACTCGAG

(c) *Methanothrix thermoacetophila sufB*C*

GTCGACAAGGAGATATACATATGTCCGACATAGCTGAGAGGGCGAGGAAAGCTCTGGGCAAGAAGGCGACATATGGCCCGGATGTGGATCTTGA
GGAGTATGAGGTCGAGGCGAAGCCACACGATGAGGTTCAGAGCATCGAGGAGGTGCCTGAGGAGGTCAGGAGGGCAGCTCTGGGTGTTGGTGTG
AGTTTGGAGGATGAGGTCGGTGGCGCATACATCCAGATGGATCAGACGCCTGTCGTCGCGAGCTCCTCCACTGAGGGGATCGAGGTGCTGGATA
TAGCCACTGCTTTGAAGAAGTATGATGGTCTCAAGGACTACTGGTGGAAGCTCGTTCCTGTGGACCAGGACAAGTACACAGCAGATGTTGCACT
CCATCCACCAGCCGGCTACTTCCTCAGGGCGAGGAGGGGCGCGAAGACGATATTTCCGCTCAAGAGCTGTCTATACCTCGGGCAGACGAATTTG
AACCAGAGAGTGCACAACATAATAATAGCGGAGGAGGGCAGCGAGCTTCACGTCATCACAGGATGCACGACGCCCAGCCATGTGCAGAGAGGCC
TGCATCTCGGCGTGTCTGAGTTCTTCATAAAGAAGGGCGCGCATGTGAGCTTCACCATGATCCACAACTGGGGGCGCGAGGTTGAGGTCCGGCC
CAGAACAGCGATATACATGGAGGGGGGGGCCACTCTATCGAACAATTACATCTGCCTGACGCCCGCGAAGACCCTGCAGACGTACCCGAAGGCG
ATACTGGATGGGCCCGGGGCGGTGGCGAGCTTCAACACCGTTCTCTATGCGAAGGAGGGGCATCTTGACACCGGAAGCGAGGTCGTGCTTAGGG
CGCCCGGATGCAGCGCGGACTCCATAACAAGAGCGGTCTCGGTCGGCGGGGAGATAATAACCAGAGGAAAGATGGTCGGCGAGGTGCCTGACGT
CAAGGCGCATCTCGAGTGCGTCGGGCTGATATTATCTGAGAAGGGTAGAATAGACTCCATACCTGAGCTCGATGGAAGATGCGGCGGCCTCGAC
ATGTCGCATGAGGCTGCAGTCGGCAAGATCTCAGAGCAGGAGCTGGAGTACCTCATGGCAAGGGGCCTGACAAAGGATCAGGCGACCGGCGTAA
TAGTCAGAGGGTTCCTGGACGTGGAGATAAAGGGCCTTCCGGAATCCCTCAGGGATGAGCTGAGGAGGATAATGCAGCTCCAGGAGCTGCACGG
AGGAGCCTAAAACGAGGGCGGGAAGCTTAAGGAGACACACTAATGTTCGGGCTGCTCGAGATCGAGGATCTGCATGTAAGCATCGGCGGCCGGA
TGGTTCTGAAGGGCATAAACCTCAAGATCAACCGGGGAGAGAACCATGCCCTCTTCGGGCCGAACGGATGCGGCAAGACCACGCTGCTGAATAC
GATAGCAGGCGTGCCCAGATATATCGTTGAAAGAGGAAGGATACTGTTCAAAGGAAATGATATTACCCATTTATCGATGGATGAACGGGCGAGG
ATGGGGATAGGGATCGCGTTTCAGAGCCCGCCTGCGATAAGGGGACTGAAGCTGAAGGATCTCATAAAGCTGTGCAACCCTGAGGTGAACCCGG
AGAAGGTGCTGAAGGAGGTGAACTTCCTGGAGTTTGCAGATCGTGAGGTCAACCGCGGATTCTCGGGAGGGGAGATCAAGAGATCCGAGTTGGC
TCAGCTGCTTGCGCAGAGACCGGAGCTTGTGATGCTGGATGAGCCTGATTCAGGAGTTGATCTGGAGAACATAAAGCTCATCGGCAGAGCGATA
AACAAGCTCACAGAGAAGGACACGAAGCCCAGCAAGAGGATCCGCTCTGGCATCGTAATAACACACTTCGGTCACATACTCGACTACATCTCTG
TGGACAAAGCTCATGTTATGATGGAGGGCACGATAGTCTGCACGGGCAGTCCGAGGGAGATACTGGAGCAGATCAAGACAAGAGGATACGAGGG
GTGTGTCAGGTGCCTATGTCCGACATAGAATTCTAGA

(d) *Methanothrix thermoacetophila iscSU**

CTCGAGAAGGAGATATACCATGAGGCGCATATACATGGACCACTCGGCCACGACCCCGGTTGCACCCGAGGTCCTGGATGCGATGCTGCCGTAC
TTCAGGGAGAAGTATGGCAACGCGTCAAGTCTCCACGAGTTCGGCCGGGAGGCGCGCGATGCTGTTGAGAGCGCGAGGGAGGAGCTCGCCTCGC
TGATCGGCGCGAGCCCTGAGGAGATATACTTCACGAGCGGGGGCACAGAGTCGGACAACCTGGCGCTCAAGGGCGTTGCGCTCAGCAGCAAGGG
AAAGCATATCATCACGACGTCGATAGAGCATCCTGCGGTTCTTGAGGTGTGCAGGTCTCTGGAGCGCATGGGCTTCGATGTCACGTGCGTTCCA
GTTGATGATCAGGGGATAGTCAATCCGGGAGAGATCGAGAGGGCGATACGCGATGATACTGTGCTCATATCCGTCATGCATGCGAACAACGAGA
TAGGTACAATACAGCCGATAAAGGAGATCGCGGAGATCGCCTCGGAGAGAGACATACTCCTGCACACGGATGCTGTCCAGACGGTTGGCAAGAT
ACCCGTGAATGTTGACCAGCTTGGTGTTGATCTGCTCTCAATATCTGCGCACAAGTTCTATGGTCCGAAGGGCGTCGGCGCGCTGTATGTCAGA
AAAGGCACGAGGATCGAGAGCATAATCCAGGGCGGTGGGCACGAGAGGGGGCTCAGGTCGGGCACAGAGAACGTACCGGGGATCGTGGGTATGG
GCAGAGCTGCAGAGCTTGCATCTGAGATGATGGATAGAGAGGCGGAGAGGCTCACAGCGCTGCGCGAGAGGCTCAAGAGTTTTGTGCTCTCGAA
CATCGACGACTCCTGGCTCAACGGCCATCCGACTAAGAGACTTCCGATCAACCTTAACTTCGGCTTCGGCAGGGTGGAGGGAGAATCGCTGCTG
CTCTACCTCGACTCAAAGGGCATAGCAGTCTCCACAGGATCAGCCTGCTCTTCAAAGAAGCTGGAGCCGAGTCACGTGCTCAGGGCCATAGGCC
TGGATCCGGTGAGATGTCATGGCTCCCTCAGGATAACGCTGGGAAGGGATAACACGCAGGAGGATGTCGATTACGTCGGTGAGTGCATACGCGA
GGCTGTGGAGCGGTTCCGCGGCATATCAGCGCTGAGGTAGAAGCTTGATATCGAATTCCTGCAGCCCGGGGGATCCACTAGTTCTAGAAATAAT
TTTGTTTAACTTTAAGAAGGAGATATACATATGATTCAGCAGACCGGCTACAGCAAAAAGGTGATGGAGCACTTCATGAACCCCAGAAATGTTG
GAGTCATAGATGATCCCGATGGATACGGCAAAGTCGGAAATCCTGTTTGTGGGGACCTCATGGAGATCTTCATAAAGGTGGGGGATGAGAAAAT
AGAGGACATCAAGTTCAGGACGTTCGGCTGCGGGGCCGCAATCGCCACGAGCAGCATGATAACGGAGATGGCTAGAGGAAAGAGCCTGGAGGAG
GCGATGAGGATAACAAGAAATGACGTGGCAGATGCTCTCGATGGCCTGCCTCCACAGAAGATGCACTGCTCCAACCTCGCGGCAGATGCTCTGC
ATGCAGCCATCAACGACTACCTCTCGAAAAAGCAAAAGCATTAAGAGCTC

(e) *Archaeoglobus fulgidus sufCB**

AAGCTTAAGGAGATATACTAATGGGCGTACTGCGCGTCGTTAACTTAGGTGTCAAGGTCGGTGAGAAGAAGATACTGAAGAACGTCAACCTGTT
CATCAAAGAGGGCGAAACCTTTGTATTATTCGGTCCTAATGGCTCAGGTAAATCGACCCTGTTGAACGCCTTAATTGGAAACCCTTCTTACAAG
GTAACCTCTGGGCGAATCTTGTTTAAGAAGGTTGATATTACTAATTTGCCCACCAACGAGCGTGTAAAGTTGGGGATCGGTATCGCCTTCCAAA
ATCCACCGAAGATAAGTGGAGTAAAGCTGATTGACGTGCTGCGATACTGCGCACGTGTTGGTGGGTACGGAGAGGACAAGATACTTGAGTTGGC
TGAGAAGATGCGAATGACTGACCATCTGTACCGTGAGATAAACATGGGGTTCTCTGGTGGAGAGGTCAAGCGTTCAGAGCTGTTACAACTCATG
CTGATGAACCCTGACCTTGTTCTTCTGGACGAACCTGACTCTGGCGTTGATTTGGAGAATGTAGCCCTTATTGGCGAAGCAATTCGTACATTGT
TAGAGCGTGATAAGGAAGAGGGTGAAAGAACCAAGTCTGGTATCATCATCACCCATCAAGGGCACATACTGGACTACGTTGACGCAGACTATGC
CGCTATCCTGTACAATGGTCGCATAGCTTGTATCGGCGAGCCTAAGGACATCCTGAACCAAATCCGAAACCACGGGTATGAGGGTTGCTTAGCG
AAGTGCCTGGCGAACATGGAGAACTCGGAATAATCTAGAGGAGAATTAATATGTTTGGTGAAATGAGTGGTGAGTACGGGGAGTTAGGGGTTGA
TAAGGAAAGATTAAAAGAGGTGGGCATCGAGCTGGACAAATCGAAGAGAAGTGGCGTTTACCTGCAAGAGGATCAAGATGCTAAAACGTTCTCC
TCATTCTTTGAGGGCGTGGAAGTAATGTCAATCAAGCAGGCAATGGAAAAGTACGACTGGGTTAAGGATTACTTCTGGAAAATCTTGCGCAAAG
ACCAAGATGAGTTCACACGTATGGCAGATACTGAAGATGTTAATGGCTACTTTATACGTTCACTTCCTGGCGCCAAAGTGGAAATTCCCGTGGA
AGCATGTTTGTACCTTAAGAAGGTGGAGAAACAGCGCGTTCATAATATTGTTATAGCTGAGGAAGGCAGCGAGCTGAATATAATCTCAGGCTGC
ACTTCCCATCCGGGCGTTGCGGGGATGCATATTGGGATAAGTGAGTTCTTCGTTAAGAAGAATGCAAAGCTCAGTTTTACGATGATCCATTCCT
GGGATACGACCATAGAGGTTCGTCCGAGAACTGCAATTAAGGTTGAAGAGGGTGGAACTTTTATTTCCAATTATATCTTACTCAATCCCGTAAA
ATTGGTGCAGACTTATCCCACCGCGTACGTAGAAAAGGACGCAACCGCCATATTTAACTCAGTAATTGTTGCCCTCGAAGGATCTGTGGTAGAC
AGCGGGAGCCGTGCTGTTCTGATGGGCGAAAACTCCCGAGCGGAAATAATTTCCCGAACGATAAGCAAGGGCGGGAAAATTATAGCGCGGGGAC
ACATCATTGGAGACGCGCCGGAAGTGAAGGGGCACCTTGAGTGTAAGGGCTTAATGCTGTCCGAATCAGGCTTGATTGACGCTATTCCCGAACT
GGAAGCCCGCTATCCCAATGTAGAGTTGTCACATGAGGCAGCAATTGGAAAGATTGCTGAAGAGGGAATCTTTTACTTAATGAGTAGAGGACTG
TCGCGCGACGAAGCCATAAGTGCGATTGTGCGCGGCTTCATGGAAATCGAGATCAAGGGTTTGCCCGAGGCTTTACAAGAGGCTATACGGCGGA
CCATCGAAATGGCAGAGCGAGACTTGCTGTAAGAGCTC

(f) *Archaeoglobus fulgidus iscS2U*2*

AAGCTTAAGGAGATATACTAATGGCCTACTTTGATTACACATCCGCGAAGCCTGTTGACGAGCGTGTATTAGAAGCCATGCTTCCTTACATGAC
TGAGAGCTTCGGGAATCCCTCAAGTGTGCACTCTTACGGATTTAAGGCGCGTGAGGCCGTTCAAGAAGCTAGAGAAAAGGTTGCAAAGTTGGTA
AATGGTGGTGGTGGCACCGTCGTTTTCACTTCGGGAGCCACTGAAGCCAACAACCTGGCCATTATCGGATACGCGATGCGTAATGCCCGGAAAG
GCAAGCACATCCTTGTCAGTGCCGTTGAGCACATGAGCGTCATTAATCCAGCCAAATTTCTGCAAAAGCAAGGATTTGAGGTGGAATATATCCC
GGTCGGTAAATACGGTGAGGTGGATGTTAGCTTCATAGACCAGAAACTTAGAGATGATACCATTTTAGTCAGTGTGCAACACGCGAACAATGAG
ATAGGGACCATTCAGCCCGTCGAAGAAATTTCCGAGGTTTTGGCTGGAAAAGCCGCACTCCACATTGATGCTACGGCCAGCGTGGGCCAAATCG
AGGTCGATGTAGAGAAAATCGGAGCCGACATGCTGACAATATCCAGCAACGACATCTACGGACCAAAAGGGGTTGGTGCCCTGTGGATAAGAAA
GGAAGCTAAGTTACAACCCGTTATATTAGGCGGTGGTCAAGAAAATGGTCTGAGATCAGGGAGCGAGAATGTGCCTTCGATCGTTGGTTTTGGT
AAGGCAGCTGAGATAACAGCGATGGAGTGGCGGGAAGAGGCGGAACGGTTACGGCGCCTTCGCGACCGTATTATCGACAATGTGCTGAAGATCG
AGGAGAGTTATCTGAACGGCCATCCGGAAAAGCGTTTGCCTAACAATGTAAATGTCCGGTTCAGTTACATCGAGGGTGAAAGCATTGTGTTATC
ACTTGACATGGCGGGGATACAAGCTAGTACCGGAAGTGCGTGTTCAAGCAAGACCCTTCAACCCAGCCATGTCCTCATGGCGTGCGGGTTGAAG
CACGAAGAAGCCCACGGTACTCTTCTTCTGACATTGGGACGATACAATACAGACGAAGATGTCGATCGACTGCTTGAGGTCCTGCCCGGCGTAA
TAGAGCGTCTGCGTAGTATGTCCCCCTTGTACCGTCGGTAATCTAGAGGAGAATTAATATGTACTCCGATAAGGTTTTCGACCACTTCCAAAAT
CCGCGTAACGTGGGAAAGATCGAGGACGCCGACGGCGTCGGGACTGTTGGAAATCCTGTCTGCGGAGACTTGATGACGATCTACATTAAGGTCA
AGGACAACCGGATTGAGGACATCAAGTTCCAAACCTTCGGATGTGCTGCGGCTATCGCTACCAGCAGCATGGCGACGGAGATGGCCAAAGGGAA
GACCATCGAGGAAGCGCTGAAGATAACACGCGACGCAGTGGCCGAGGCTTTGGGTGGTCTGCCAAAGCAAAAGATGCATTGTTCTAATTTAGCT
GCCGATGCTTTACGGCGCGCTATAGTGGACTATTTTCGCAAGAATGGCAAGATAGACAAAATTAAGGAACTGGGACTGGAAAAGGAGTTAGAAA
AGATGGAAAAGGGTGAAATGGATGACCACGGCGAGTACTGCGAGGCCTAAGAGCTC

**Legends to supplementary figures**

**Figure S1.** Comparison of amino acid sequences. Multi-sequence alignments are provided for IscS, NifS and SufS (a), for IscU, NifU, IscU* and SufU (b), for SufC (c), and for SufB, SufB* and SufD (d). The alignments were performed using Clustal Omega (https://www.ebi.ac.uk/Tools/msa/clustalo/), and the figure was generated using ESPript (http://espript.ibcp.fr/ESPript/ESPript/). Abbreviations: Af, *Archaeoglobus fulgidus*; Bs, *Bacillus subtilis*; Cp, *Clostridium perfringens*; Ec, *Escherichia coli*; Hp, *Helicobacter pylori*; Mj, *Methanocaldococcus jannaschii*; Mt, *Methanothrix thermoacetophila*; and Ta, *Thermoplasma acidophilum*.

**Figure S2.** Complementation tests of *E. coli* UT109 by *sufB*C* and/or *apbC* from *M. jannaschii.* The UT109 cells harboring pUMV22 $Sp^r$ were transformed with the compatible plasmids pBBR-*Mj sufB*C* and/or pRK-*Mj apbC.* Cells were grown on LB-glucose plates supplemented with 5 mM $Na_2S$ (without MVA) at 37˚C for 72 hr under anaerobic conditions.

**Figure S3.** Exogenously supplied $Na_2S$ could not substitute for the roles of *E. coli* SUF components. UT109 cells harboring the plasmid expressing *E. coli* SUF machinery each lacking a single SUF component (or SufSE) were grown on a LB-glucose plate supplemented with 5 mM $Na_2S$ (without MVA) at 37˚C for 72 hr under anaerobic conditions.

**Figure S4.** SDS-PAGE analysis of the purified components of the *Mj* SufB*C complex and its variant forms.

**Figure S5.** The loci of amino acid substitutions that allowed *Mj* SufB*1 and *Mj* SufB*2 to work asymmetrically. In the AlphaFold2-predicted structure of the *M. jannaschii* SufB* homodimer, the positions of suppressor mutations are indicated by the wild-type side chains in sticks. The essential residues of *Mj* SufB* (Cys252, His279 and Glu280) are depicted as spheres.

**Figure S6.** Phylogenetic tree of SufB*. The evolutionary history of the 137 amino acid sequences of the SufB* subfamily was inferred using the maximum-likelihood method in the MEGA X software. Colors of branches correspond to archaea (red), clostridia (green), delta-proteobacteria (blue), and other bacteria (black). Co-occurrence of the genes for IscU* in the respective genomes is indicated by dots.

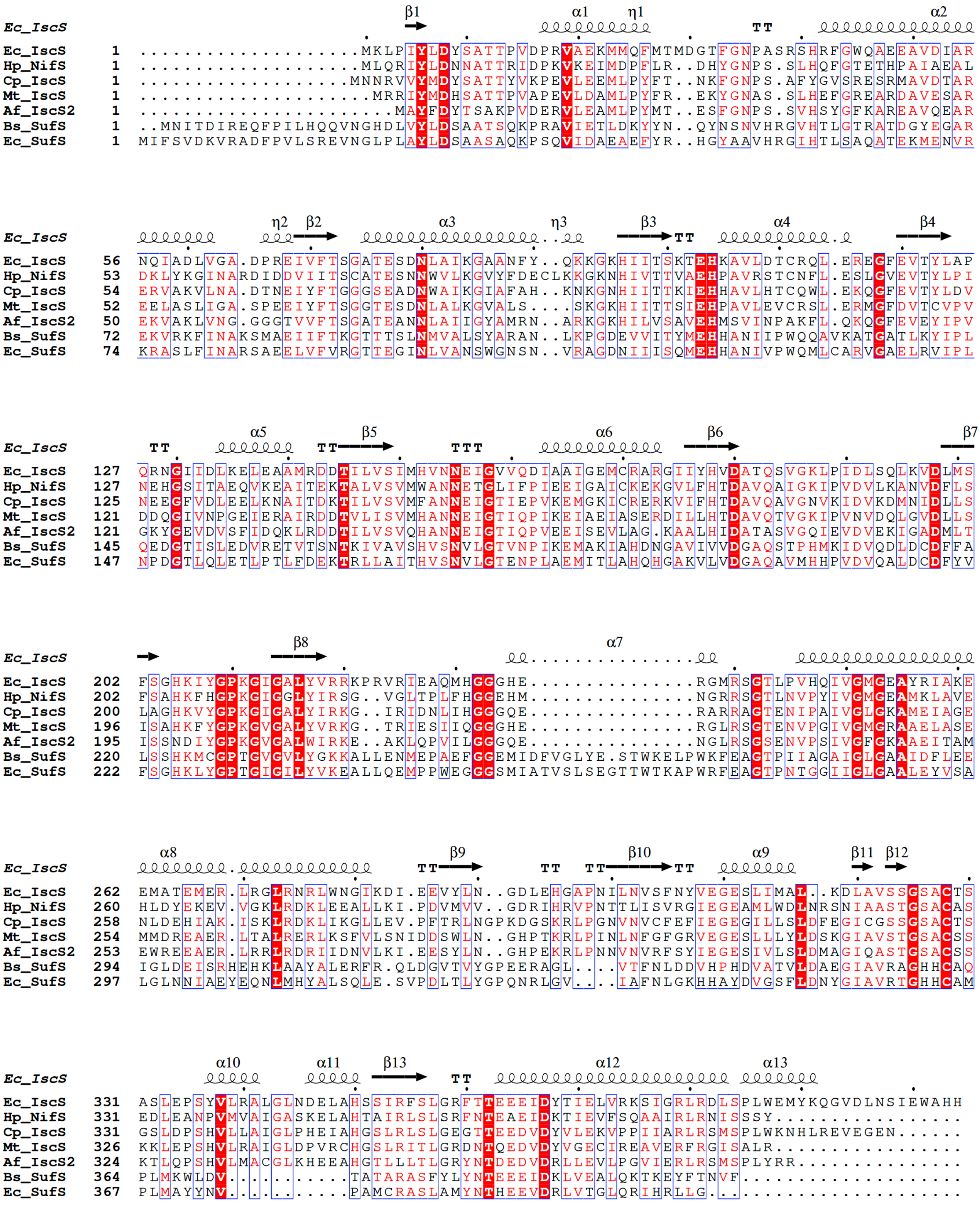


Figure S1a

```
                                 α1                 TT           β1        β2
Ec_IscU
Ec_IscU     1  ............MAYSEKVIDHYENPRNVGSFDNNDEN.....VGSGMVGAPACGDVMKLQIKVND.E
Hp_NifU     1  MAKHDLVGSALWDAYSKEVQRRMDNPTHLGVITEEQAKAKNAKLIVADYGAEACGDAVRLYWLVDEST
Cp_IscU*    1  ............MIYSEKVMDHFKNPRNVGEIKDA........NGIGEVGNAKCGDIMRIYLKIEN.N
Mt_IscU*    1  ........MIQQTGYSKKVMEHFMNPRNVGVIDDP........DGYGKVGNPVCGDLMEIFIKVGD.E
Af_IscU*    1  .............MYSDKVFDHFQNPRNVGKIEDA........DGVGTVGNPVCGDLMTIYIKVKD.N
Bs_SufU     1  ....MSFNANLDTLYRQVIMDHYKNPRNKGVLNDS.........IVVDMNNPTCGDRIRLTMKLDG..

                     β3               α2            α3          α4
Ec_IscU
Ec_IscU    51  GIIEDARFKTYGCGSAIASSSLVTEWVKGKSLDEAQAIKNTDIAEE..........L.........EL
Hp_NifU    69  DTIIDAKFKSFGCGTAIASSDMMVELCLNKRVQDAVKITNLDVERG..........LRDDP..DTPAV
Cp_IscU*   48  .IVEDVKFETYGCGSAIASSSMATELIKGKTVEEAWELSNKAVAEA..........LD........GL
Mt_IscU*   52  .KIEDIKFRTFGCGAAIATSSMITEMARGKSLEEAMRITRNDVADA..........LD........GL
Af_IscU*   47  .RIEDIKFQTFGCAAAIATSSMATEMAKGKTIEEALKITRDAVAEA..........LG........GL
Bs_SufU    54  DIVEDAKFEGEGCSISMASASMMTQAIKGKDIETALSMSKIFSDMMQGKEYDDSIDLGDIEALQGVSK

                 η1       α5
Ec_IscU
Ec_IscU   100  PPVKIHCSILAEDAIKAAIADYKSKREAK.......................................
Hp_NifU   125  PGQKMHCSVMAYDVIKKAAGMYLGKNAEDFEEEIIVCECARVSLGTIKEVIRLNDLKSVEEITNYTKA
Cp_IscU*   97  PPVKMHCSVLAEEAIHKAINDYRQRQGLEPWDMK.........E.HSHDIHE.......HVHGQ....
Mt_IscU*  101  PPQKMHCSNLAADALHAAINDYLSKKQKH.......................................
Af_IscU*   96  PKQKMHCSNLAADALRRAIVDYFRKNGKIDKIKE.........LGLEKELEK.......MEKGEMDDH
Bs_SufU   122  FPARIKCATLSWKALEKGVAKEEGGN..........................................

Ec_IscU
Ec_IscU        ....................................................................
Hp_NifU   193  GAFCKSCVRPGGHEKRDYYLVDILKEVREEMEAEKLKAAANKSQNGELAFREMTMVQKIKAVDKVIDE
Cp_IscU*  144  ..E.................................................................
Mt_IscU*       ....................................................................
Af_IscU*  148  GEYCEA..............................................................
Bs_SufU        ....................................................................

Ec_IscU
Ec_IscU        ..................................................................
Hp_NifU   261  NIRAMLMMDGGDLEILDIKESDDYIDVYIRYMGACDGCMSATTGTLFAIENALQELLDRSIRVLPI
Cp_IscU*       ..................................................................
Mt_IscU*       ..................................................................
Af_IscU*       ..................................................................
Bs_SufU        ..................................................................
```

Figure S1b

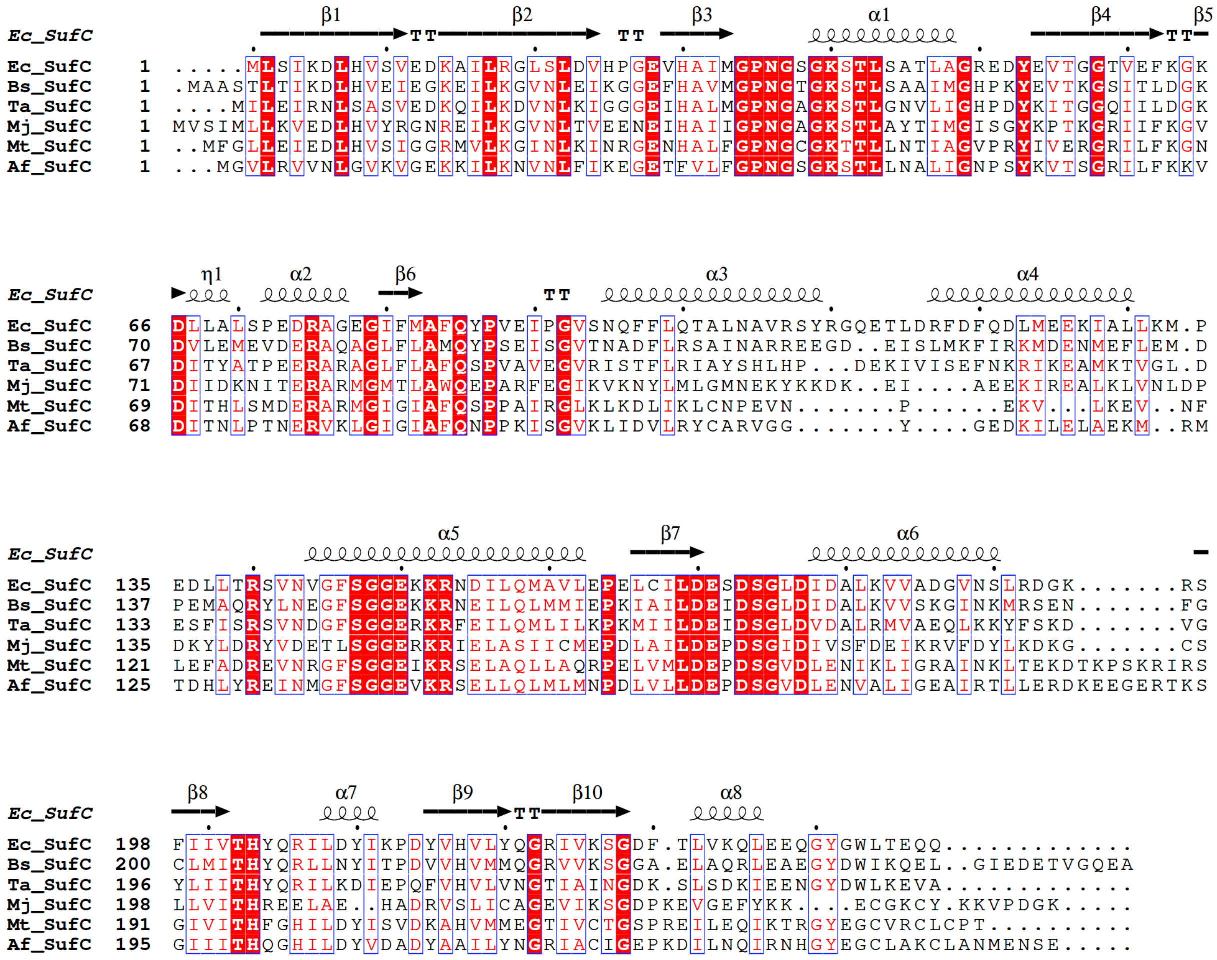
β1 β2 β3 α1 β4 β5
Ec_SufC
Ec_SufC 1 .....MLSIKDLHVSVEDKAILRGLSLDVHPGEVHAIMGPNGSGKSTLSATLAGREDYEVTGGTVEFKGK
Bs_SufC 1 .MAASTLTIKDLHVEIEGKEILKGVNLEIKGGEFHAVMGPNGTGKSTLSAAIMGHPKYEVTKGSITLDGK
Ta_SufC 1 ....MILEIRNLSASVEDKQILKDVNLKIGGGEIHALMGPNGAGKSTLGNVLIGHPDYKITGGQIILDGK
Mj_SufC 1 MVSIMLLKVEDLHVYRGNREILKGVNLTVEENEIHAIIGPNGAGKSTLAYTIMGISGYKPTKGRIIFKGV
Mt_SufC 1 ..MFGLLEIEDLHVSIGGRMVLKGINLKINRGENHALFGPNGCGKTTLLNTIAGVPRYIVERGRILFKGN
Af_SufC 1 ...MGVLRVVNLGVKVGEKKILKNVNLFIKEGETFVLFGPNGSGKSTLLNALIGNPSYKVTSGRILFKKV
η1 α2 β6 α3 α4
Ec_SufC
Ec_SufC 66 DLLALSPEDRAGEGIFMAFQYPVEIPGVSNQFFLQTALNAVRSYRGQETLDRFDFQDLMEEKIALLKM.P
Bs_SufC 70 DVLEMEVDERAQAGLFLAMQYPSEISGVTNADFLRSAINARREEGD..EISLMKFIRKMDENMEFLEM.D
Ta_SufC 67 DITYATPEERARAGLFLAFQSPVAVEGVRISTFLRIAYSHLHP...DEKIVISEFNKRIKEAMKTVGL.D
Mj_SufC 71 DIIDKNITERARMGMTLAWQEPARFEGIKVKNYLMLGMNEKYKKDK..EI....AEEKIREALKLVNLDP
Mt_SufC 69 DITHLSMDERARMGIGIAFQSPPAIRGLKLKDLIKLCNPEVN.......P......EKV...LKEV..NF
Af_SufC 68 DITNLPTNERVKLGIGIAFQNPPKISGVKLIDVLRYCARVGG.......Y....GEDKILELAEKM..RM
α5 β7 α6
Ec_SufC
Ec_SufC 135 EDLLTRSVNVGFSGGEKKRNDILQMAVLEPELCILDESDSGLDIDALKVVADGVNSLRDGK.......RS
Bs_SufC 137 PEMAQRYLNEGFSGGEKKRNEILQLMMIEPKIAILDEIDSGLDIDALKVVSKGINKMRSEN.......FG
Ta_SufC 133 ESFISRSVNDGFSGGERKRFEILQMLILKPKMIILDEIDSGLDVDALRMVAEQLKKYFSKD.......VG
Mj_SufC 135 DKYLDRYVDETLSGGERKRIELASIICMEPDLAILDEPDSGIDIVSFDEIKRVFDYLKDKG.......CS
Mt_SufC 121 LEFADREVNRGFSGGEIKRSELAQLLAQRPELVMLDEPDSGVDLENIKLIGRAINKLTEKDTKPSKRIRS
Af_SufC 125 TDHLYREINMGFSGGEVKRSELLQLMLMNPDLVLLDEPDSGVDLENVALIGEAIRTLLERDKEEGERTKS
β8 α7 β9 β10 α8
Ec_SufC
Ec_SufC 198 FIIVTHYQRILDYIKPDYVHVLYQGRIVKSGDF.TLVKQLEEQGYGWLTEQQ.............
Bs_SufC 200 CLMITHYQRLLNYITPDVVHVMMQGRVVKSGGA.ELAQRLEAEGYDWIKQEL..GIEDETVGQEA
Ta_SufC 196 YLIITHYQRILKDIEPQFVHVLVNGTIAINGDK.SLSDKIEENGYDWLKEVA.............
Mj_SufC 198 LLVITHREELAE..HADRVSLICAGEVIKSGDPKEVGEFYKK....ECGKCY.KKVPDGK.....
Mt_SufC 191 GIVITHFGHILDYISVDKAHVMMEGTIVCTGSPREILEQIKTRGYEGCVRCLCPT..........
Af_SufC 195 GIIITHQGHILDYVDADYAAILYNGRIACIGEPKDILNQIRNHGYEGCLAKCLANMENSE.....

Figure S1c

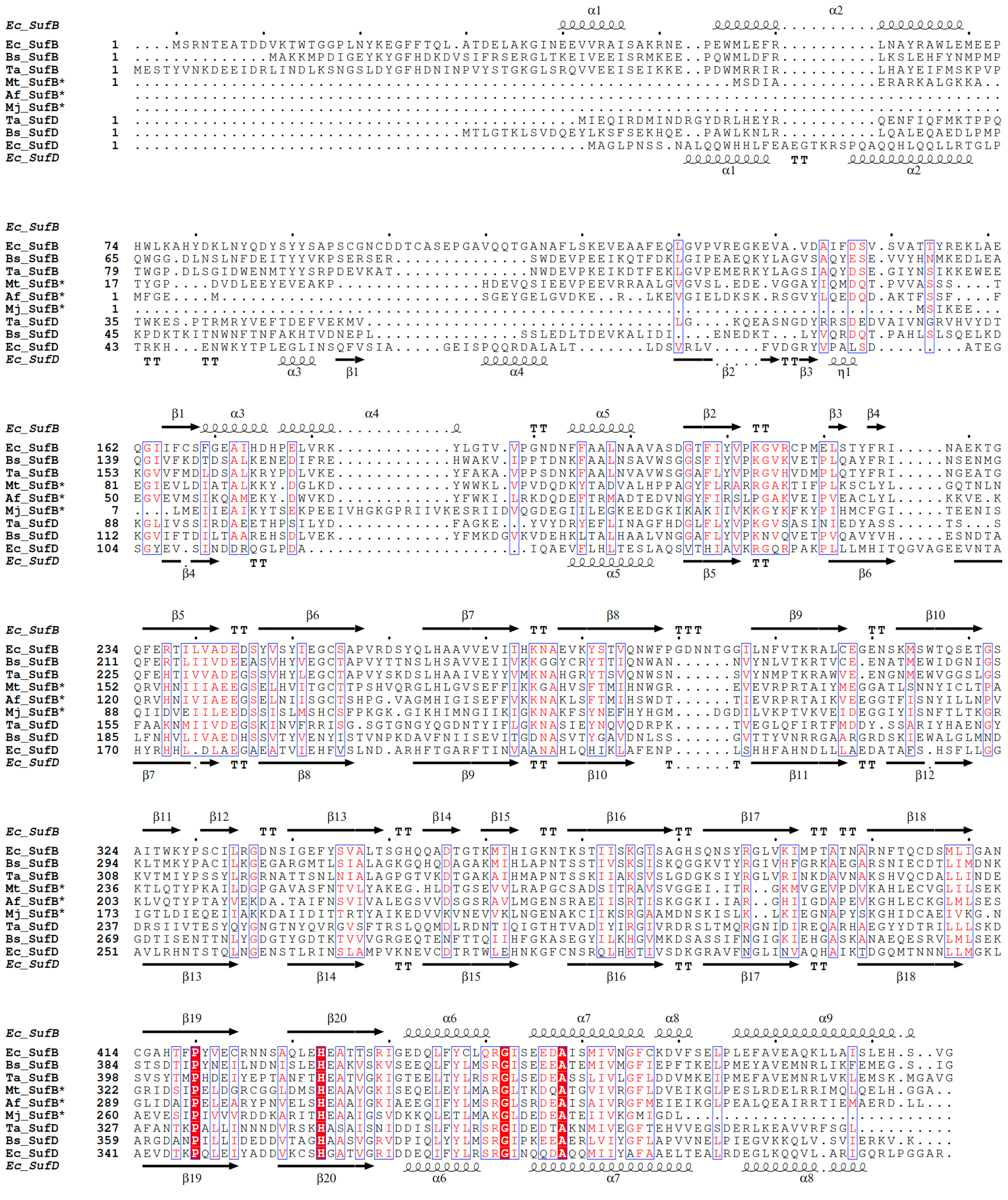

Figure S1d

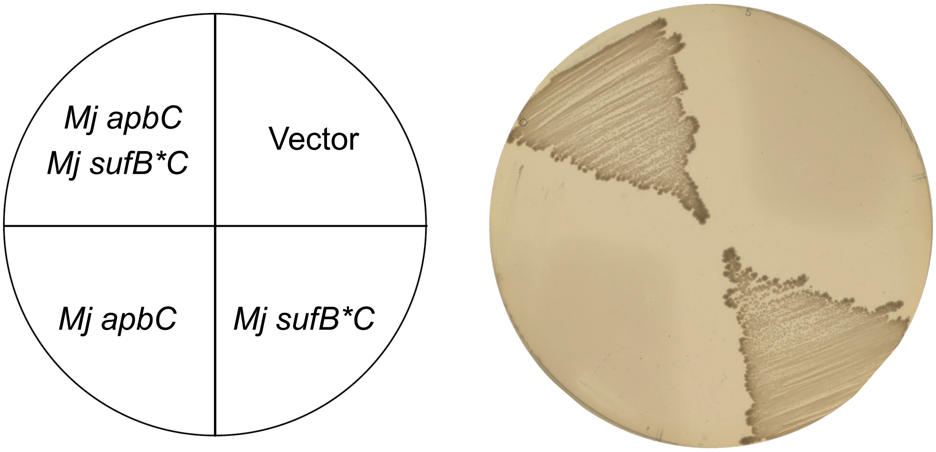


Figure S2

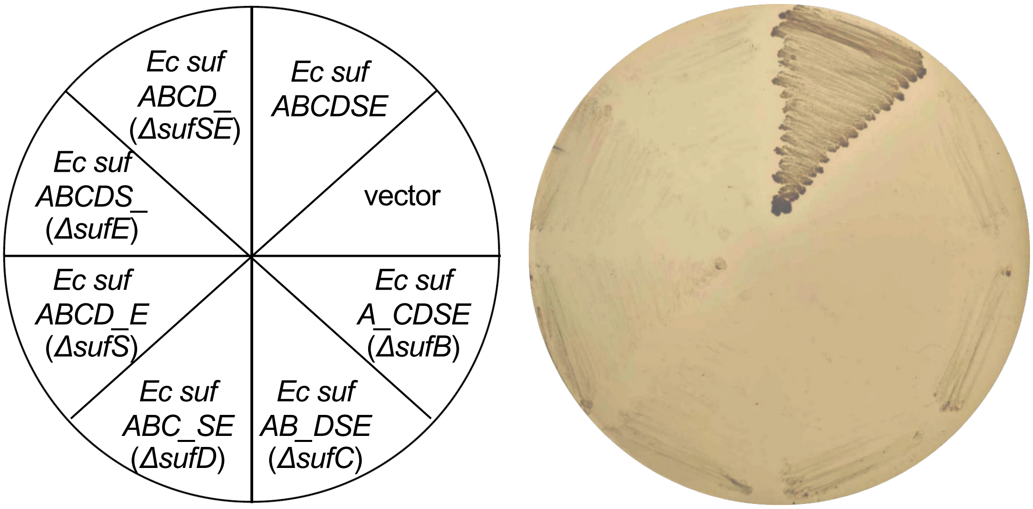
Ec suf
ABCDSE
vector
Ec suf
A_CDSE
(ΔsufB)
Ec suf
AB_DSE
(ΔsufC)
Ec suf
ABC_SE
(ΔsufD)
Ec suf
ABCD_E
(ΔsufS)
Ec suf
ABCDS_
(ΔsufE)
Ec suf
ABCD_
(ΔsufSE)

Figure S3

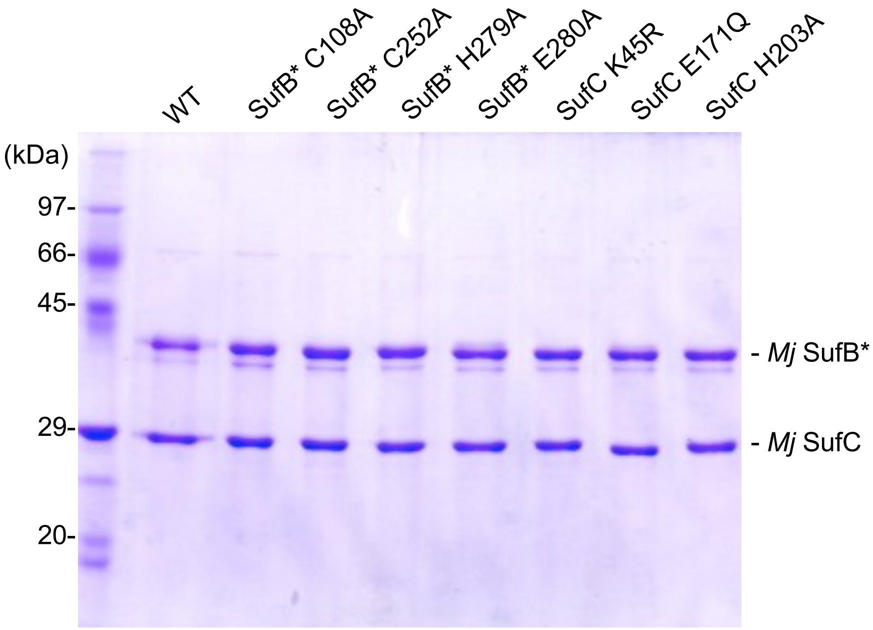
WT
SufB* C108A
SufB* C252A
SufB* H279A
SufB* E280A
SufC K45R
SufC E171Q
SufC H203A
(kDa)
97-
66-
45-
29-
20-
- Mj SufB*
- Mj SufC

Figure S4

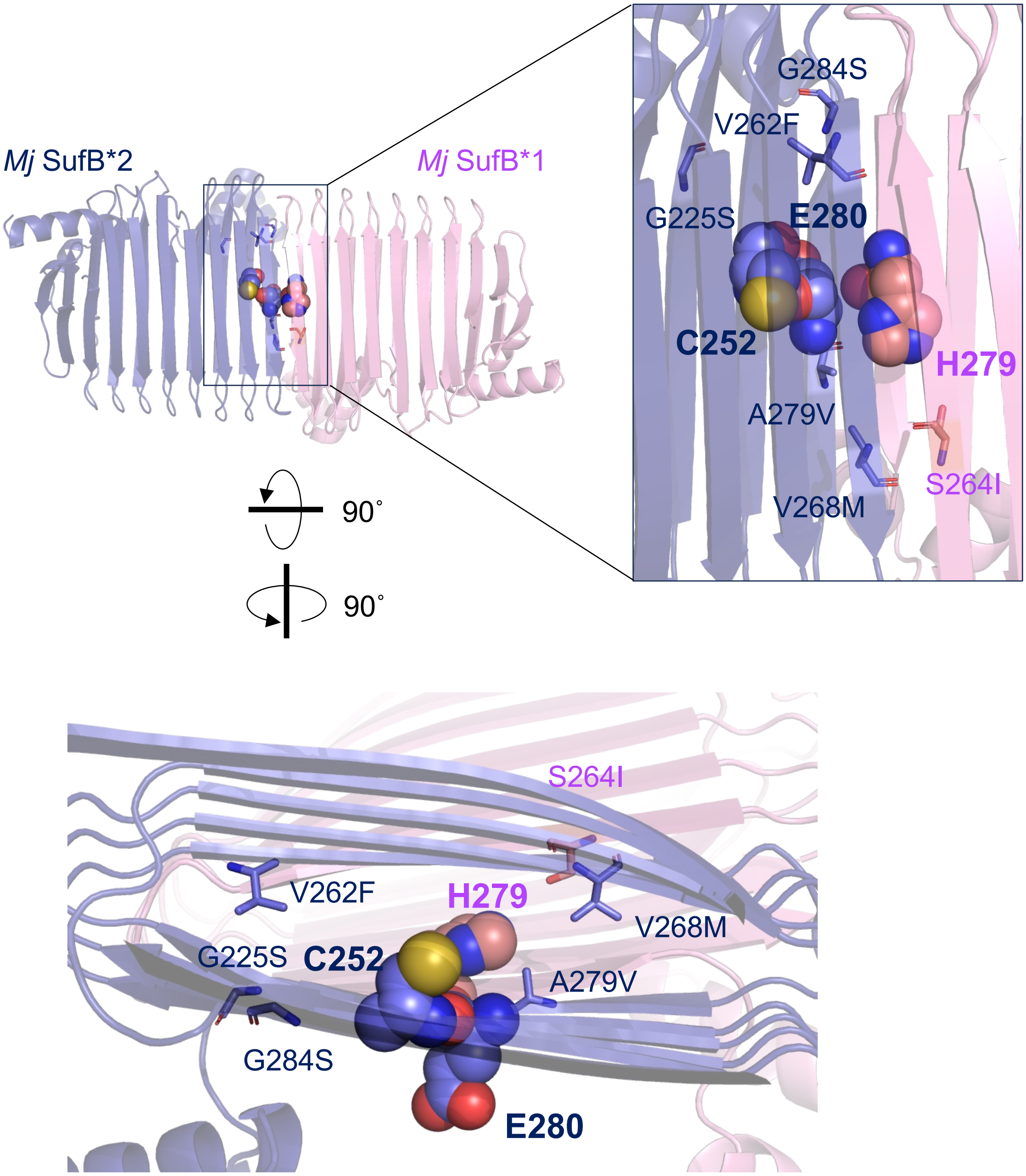

*Mj* SufB*2
*Mj* SufB*1
G284S
V262F
G225S
E280
C252
H279
A279V
S264I
V268M
90˚
90˚
S264I
V262F
H279
V268M
G225S
C252
A279V
G284S
E280


Figure S5

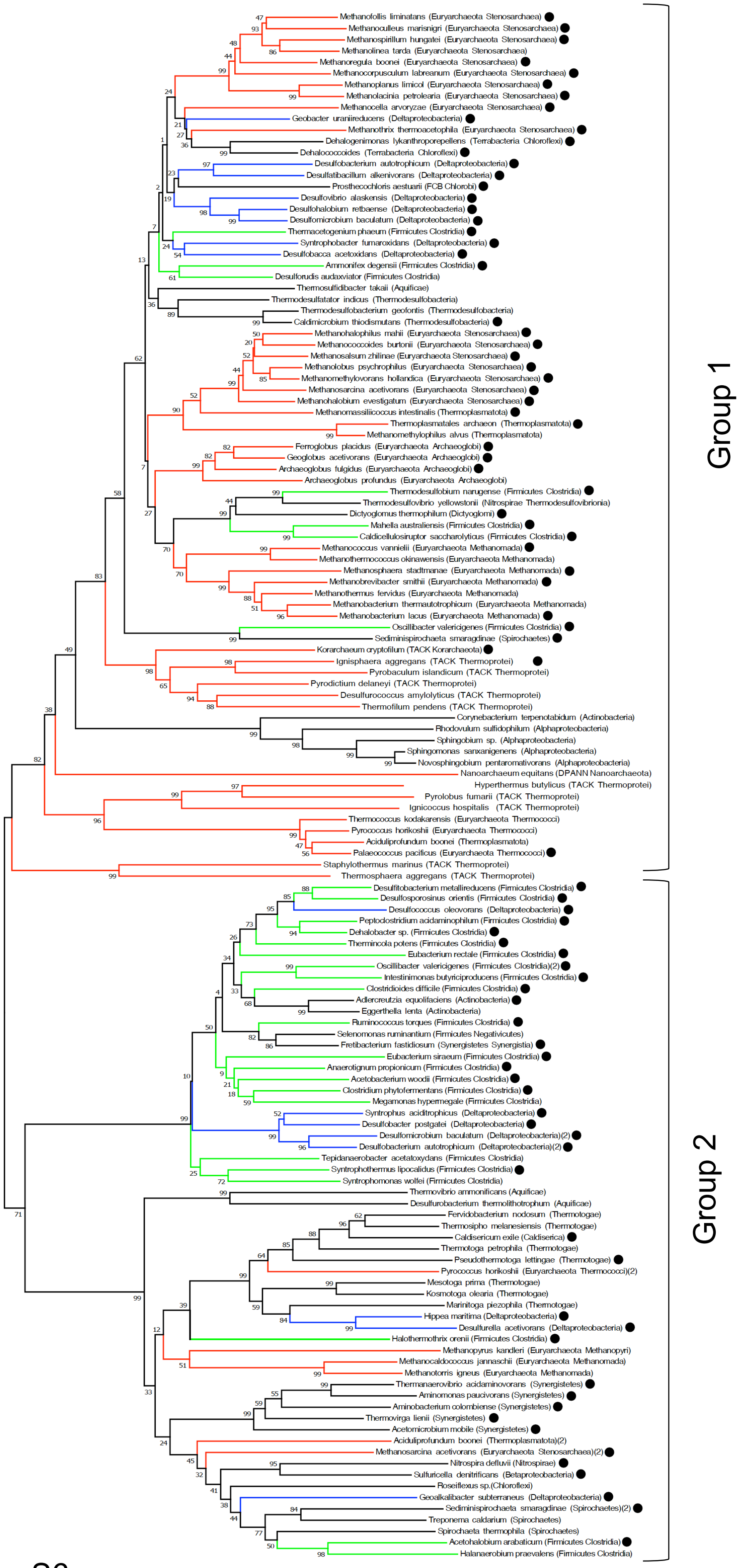

Methanofollis liminatans (Euryarchaeota Stenosarchaea)
Methanoculleus marisnigri (Euryarchaeota Stenosarchaea)
Methanospirillum hungatei (Euryarchaeota Stenosarchaea)
Methanolinea tarda (Euryarchaeota Stenosarchaea)
Methanoregula boonei (Euryarchaeota Stenosarchaea)
Methanocorpusculum labreanum (Euryarchaeota Stenosarchaea)
Methanoplanus limicol (Euryarchaeota Stenosarchaea)
Methanolacinia petrolearia (Euryarchaeota Stenosarchaea)
Methanocella arvoryzae (Euryarchaeota Stenosarchaea)
Geobacter uraniireducens (Deltaproteobacteria)
Methanothrix thermoacetophila (Euryarchaeota Stenosarchaea)
Dehalogenimonas lykanthroporepellens (Terrabacteria Chloroflexi)
Dehalococcoides (Terrabacteria Chloroflexi)
Desulfobacterium autotrophicum (Deltaproteobacteria)
Desulfatibacillum alkenivorans (Deltaproteobacteria)
Prosthecochloris aestuarii (FCB Chlorobi)
Desulfovibrio alaskensis (Deltaproteobacteria)
Desulfohalobium retbaense (Deltaproteobacteria)
Desulfomicrobium baculatum (Deltaproteobacteria)
Thermacetogenium phaeum (Firmicutes Clostridia)
Syntrophobacter fumaroxidans (Deltaproteobacteria)
Desulfobacca acetoxidans (Deltaproteobacteria)
Ammonifex degensii (Firmicutes Clostridia)
Desulforudis audaxviator (Firmicutes Clostridia)
Thermosulfidibacter takaii (Aquificae)
Thermodesulfatator indicus (Thermodesulfobacteria)
Thermodesulfobacterium geofontis (Thermodesulfobacteria)
Caldimicrobium thiodismutans (Thermodesulfobacteria)
Methanohalophilus mahii (Euryarchaeota Stenosarchaea)
Methanococcoides burtonii (Euryarchaeota Stenosarchaea)
Methanosalsum zhilinae (Euryarchaeota Stenosarchaea)
Methanolobus psychrophilus (Euryarchaeota Stenosarchaea)
Methanomethylovorans hollandica (Euryarchaeota Stenosarchaea)
Methanosarcina acetivorans (Euryarchaeota Stenosarchaea)
Methanohalobium evestigatum (Euryarchaeota Stenosarchaea)
Methanomassiliicoccus intestinalis (Thermoplasmatota)
Thermoplasmatales archaeon (Thermoplasmatota)
Methanomethylophilus alvus (Thermoplasmatota)
Ferroglobus placidus (Euryarchaeota Archaeoglobi)
Geoglobus acetivorans (Euryarchaeota Archaeoglobi)
Archaeoglobus fulgidus (Euryarchaeota Archaeoglobi)
Archaeoglobus profundus (Euryarchaeota Archaeoglobi)
Thermodesulfobium narugense (Firmicutes Clostridia)
Thermodesulfovibrio yellowstonii (Nitrospirae Thermodesulfovibrionia)
Dictyoglomus thermophilum (Dictyoglomi)
Mahella australiensis (Firmicutes Clostridia)
Caldicellulosiruptor saccharolyticus (Firmicutes Clostridia)
Methanococcus vannielii (Euryarchaeota Methanomada)
Methanothermococcus okinawensis (Euryarchaeota Methanomada)
Methanosphaera stadtmanae (Euryarchaeota Methanomada)
Methanobrevibacter smithii (Euryarchaeota Methanomada)
Methanothermus fervidus (Euryarchaeota Methanomada)
Methanobacterium thermautotrophicum (Euryarchaeota Methanomada)
Methanobacterium lacus (Euryarchaeota Methanomada)
Oscillibacter valericigenes (Firmicutes Clostridia)
Sediminispirochaeta smaragdinae (Spirochaetes)
Korarchaeum cryptofilum (TACK Korarchaeota)
Ignisphaera aggregans (TACK Thermoprotei)
Pyrobaculum islandicum (TACK Thermoprotei)
Pyrodictium delaneyi (TACK Thermoprotei)
Desulfurococcus amylolyticus (TACK Thermoprotei)
Thermofilum pendens (TACK Thermoprotei)
Corynebacterium terpenotabidum (Actinobacteria)
Rhodovulum sulfidophilum (Alphaproteobacteria)
Sphingobium sp. (Alphaproteobacteria)
Sphingomonas sanxanigenens (Alphaproteobacteria)
Novosphingobium pentaromativorans (Alphaproteobacteria)
Nanoarchaeum equitans (DPANN Nanoarchaeota)
Hyperthermus butylicus (TACK Thermoprotei)
Pyrolobus fumarii (TACK Thermoprotei)
Ignicoccus hospitalis (TACK Thermoprotei)
Thermococcus kodakarensis (Euryarchaeota Thermococci)
Pyrococcus horikoshii (Euryarchaeota Thermococci)
Aciduliprofundum boonei (Thermoplasmatota)
Palaeococcus pacificus (Euryarchaeota Thermococci)
Staphylothermus marinus (TACK Thermoprotei)
Thermosphaera aggregans (TACK Thermoprotei)
Desulfitobacterium metallireducens (Firmicutes Clostridia)
Desulfosporosinus orientis (Firmicutes Clostridia)
Desulfococcus oleovorans (Deltaproteobacteria)
Peptoclostridium acidaminophilum (Firmicutes Clostridia)
Dehalobacter sp. (Firmicutes Clostridia)
Thermincola potens (Firmicutes Clostridia)
Eubacterium rectale (Firmicutes Clostridia)
Oscillibacter valericigenes (Firmicutes Clostridia)(2)
Intestinimonas butyriciproducens (Firmicutes Clostridia)
Clostridioides difficile (Firmicutes Clostridia)
Adlercreutzia equolifaciens (Actinobacteria)
Eggerthella lenta (Actinobacteria)
Ruminococcus torques (Firmicutes Clostridia)
Selenomonas ruminantium (Firmicutes Negativicutes)
Fretibacterium fastidiosum (Synergistetes Synergistia)
Eubacterium siraeum (Firmicutes Clostridia)
Anaerotignum propionicum (Firmicutes Clostridia)
Acetobacterium woodii (Firmicutes Clostridia)
Clostridium phytofermentans (Firmicutes Clostridia)
Megamonas hypermegale (Firmicutes Clostridia)
Syntrophus aciditrophicus (Deltaproteobacteria)
Desulfobacter postgatei (Deltaproteobacteria)
Desulfomicrobium baculatum (Deltaproteobacteria)(2)
Desulfobacterium autotrophicum (Deltaproteobacteria)(2)
Tepidanaerobacter acetatoxydans (Firmicutes Clostridia)
Syntrophothermus lipocalidus (Firmicutes Clostridia)
Syntrophomonas wolfei (Firmicutes Clostridia)
Thermovibrio ammonificans (Aquificae)
Desulfurobacterium thermolithotrophum (Aquificae)
Fervidobacterium nodosum (Thermotogae)
Thermosipho melanesiensis (Thermotogae)
Caldisericum exile (Caldiserica)
Thermotoga petrophila (Thermotogae)
Pseudothermotoga lettingae (Thermotogae)
Pyrococcus horikoshii (Euryarchaeota Thermococci)(2)
Mesotoga prima (Thermotogae)
Kosmotoga olearia (Thermotogae)
Marinitoga piezophila (Thermotogae)
Hippea maritima (Deltaproteobacteria)
Desulfurella acetivorans (Deltaproteobacteria)
Halothermothrix orenii (Firmicutes Clostridia)
Methanopyrus kandleri (Euryarchaeota Methanopyri)
Methanocaldococcus jannaschii (Euryarchaeota Methanomada)
Methanotorris igneus (Euryarchaeota Methanomada)
Thermanaerovibrio acidaminovorans (Synergistetes)
Aminomonas paucivorans (Synergistetes)
Aminobacterium colombiense (Synergistetes)
Thermovirga lienii (Synergistetes)
Acetomicrobium mobile (Synergistetes)
Aciduliprofundum boonei (Thermoplasmatota)(2)
Methanosarcina acetivorans (Euryarchaeota Stenosarchaea)(2)
Nitrospira defluvii (Nitrospirae)
Sulfuricella denitrificans (Betaproteobacteria)
Roseiflexus sp.(Chloroflexi)
Geoalkalibacter subterraneus (Deltaproteobacteria)
Sediminispirochaeta smaragdinae (Spirochaetes)(2)
Treponema caldarium (Spirochaetes)
Spirochaeta thermophila (Spirochaetes)
Acetohalobium arabaticum (Firmicutes Clostridia)
Halanaerobium praevalens (Firmicutes Clostridia)
Group 1
Group 2


Figure S6
